# Structural basis of EHEP-mediated offense against phlorotannin-induced defense from brown algae to protect *aku*BGL activity

**DOI:** 10.1101/2023.06.04.543649

**Authors:** Xiaomei Sun, Yuxin Ye, Naofumi Sakurai, Hang Wang, Koji Kato, Jian Yu, Keizo Yuasa, Akihiko Tsuji, Min Yao

**Affiliations:** Faculty of Advanced Life Science, Hokkaido University, Sapporo, Japan; Graduate School of Bioscience and Bioindustry, Tokushima University, Tokushima, Japan

## Abstract

The defensive-offensive associations between algae and herbivores determine marine ecology. Brown algae utilize phlorotannin as their chemical defense against the predator *Aplysia kurodai*, which uses β-glucosidase (*aku*BGL) to digest the laminarin in algae to glucose. Moreover, *A. kurodai* employs *Eisenia* hydrolysis-enhancing protein (EHEP) as an offense to protect *aku*BGL activity from phlorotannin inhibition by precipitating phlorotannin. To underpin the molecular mechanism of this digestive-defensive-offensive system, we determined the structures of apo and tannic-acid (TNA, a phlorotannin-analog) bound form of EHEP, as well as apo *aku*BGL. EHEP consisted of three peritrophin-A domains formed in a triangle and bound TNA in the center without significant conformational changes. Structural comparison between EHEP and EHEP–TNA led us to find that EHEP can be resolubilized from phlorotannin-precipitation at an alkaline pH, which reflects a requirement in the digestive tract. *aku*BGL contained two GH1 domains, only one of which conserved the active site. Combining docking analysis, we propose the mechanisms by which phlorotannin inhibits *aku*BGL by occupying the substrate-binding pocket, and EHEP protects *aku*BGL against the inhibition by binding with phlorotannin to free the *aku*BGL pocket.

## Introduction

Over millions of years of evolution, predators have successfully coevolved with their prey to maintain an ecological balance ^1^. In marine habitats, interactions between algae and marine herbivores dominate marine ecosystems. Most algae are consumed by marine herbivores ^2^. They produce secondary metabolites as a chemical defense to protect themselves against predators. In brown algae *Eisenia bicyclis*, laminarin is a major storage carbohydrate, constituting 20%–30% of algae dry weight. The sea hare *Aplysia kurodai*, a marine gastropod, preferentially feeds on the *E. bicyclis* with its 110 and 210 kDa β-glucosidases (*aku*BGLs), hydrolyzing the laminarin and releasing large amounts of glucose. Interestingly, such a feeding strategy has attracted attention for producing glucose as a renewable biofuel source ^3^. However, to protect themselves against predators, brown algae produce phlorotannin, a secondary metabolite, thereby reducing the digestion by *A. kurodai* by inhibiting the hydrolytic activity of *aku*BGLs. As the 110 kDa *aku*BGL is more sensitive to phlorotannin than the 210 kDa BGL ^4^, we focused on the 110 kDa *aku*BGL in this study (hereafter, *aku*BGL refers to 110 kDa *aku*BGL).

To counteract the antipredator adaptations of algae, herbivores use diverse approaches, such as detoxification, neutralization, defense suppression, and physiological adaptations ^5^. *A. kurodai* inhibits the phlorotannin-defense of brown algae through *Eisenia* hydrolysis-enhancing protein (EHEP), a protein from their digestive system that protects *aku*BGL activity from phlorotannin inhibition ^4^. Previous studies have shown that incubating *E. bicyclis* with *aku*BGL in the presence of EHEP results in increased glucose production because EHEP binds to phlorotannin and forms an insoluble complex ^4^.

The *aku*BGL–phlorotannin/laminarin–EHEP system exemplifies the digestion process of *A. kurodai* as well as the defense and antidefense strategies between *E. bicyclis* and *A. kurodai*. Although the defense/antidefense strategy has been established, the detailed molecular mechanism of this interplay remains unknown. Further, phlorotannin inhibition hinders the potential application of brown algae as feedstocks for enzymatically producing biofuel from laminarin. Thus, understanding the underlying molecular mechanisms will be beneficial for the application of this system in the biofuel industry.

Despite the potential use of laminarin hydrolytic enzymes in the biofuel industry, only a few BGLs of glycoside hydrolases belonging to the GH3 and GH1 family are known to hydrolyze laminarin (e.g., *Talaromyces amestolkiae* BGL^6^, *Ustilago esculenta* BGL ^7^, and V*ibrio campbellii* BGL ^8^ from the GH3 family and *Saccharophagus degradans* 2-40^T^ BGL ^9^ from the GH1 family). GH3 is a multidomain enzyme family characterized by N-terminal (β/α)_8_ (NTD) and C-terminal (β/α)_6_ (CTD) domains, with or without auxiliary domains ^10^; the nucleophile aspartate and the acid/base glutamate residues exist in the NTD and CTD, respectively. In contrast, the members of the GH1 family generally share a single (β/α)_8_-fold domain (hereafter referred to as GH1 domain [GH1D]), and the two glutamic acid catalytic residues are located in the carboxyl termini of β-strands 4 and 7. Therefore, the two families may use different substrate-recognition and catalytic mechanisms for laminarin. Intriguingly, although *aku*BGL possesses laminarin hydrolytic activity and belongs to the GH1 family, its molecular weight is considerably greater than that of other GH1 members. Sequence analysis has indicated that *aku*BGL consists of ≥2 GH1Ds. Because no structural information of BGL active on polysaccharides is available, the catalytic mechanism toward laminarin is unclear.

There is limited information on EHEP, a novel cysteine-rich protein (8.2% of the amino acid content), because no structural or functional homologous protein exists in other organisms. EHEP was predicted to consist of three peritrophin-A domains (PADs) with a cysteine-spacing pattern of CX_15_CX_5_CX_9_CX_12_CX_5-9_C. The PADs consist of peritrophic matrix proteins, which have been proposed to play an important role in detoxifying ingested xenobiotics ^11^. For instance, *Aedes aegypti* intestinal mucin 1 (*Ae*IMUC1) consists of a signal peptide followed by three PADs with an intervening mucin-like domain; its expression is induced by blood feeding. *Ae*IMUC1-mediated blood detoxification during digestion is completed by binding to toxic heme molecules^12^. Despite the similar domain organization of EHEP and *Ae*IMUC1, their function and binding partner are completely different, implying their different characteristics. However, the characteristics of the EHEP-phlorotannin insoluble complex remain unknown; moreover, it remains unclear why and how EHEP protects *aku*BGL from phlorotannin inhibition.

In this study, we determined the structures of apo and tannic acid (TNA, phlorotannin-analog) bound-form of EHEP (EHEP, EHEP–TNA), as well as *aku*BGL; all isolated from *A. kurodai*. The structure of EHEP consists of three PADs arranged in a triangle shape, with TNA bound at the surface of the triangle center. A structural comparison of EHEP and EHEP–TNA revealed no significant changes in conformation upon TNA binding, implying that EHEP maintains its structure when precipitated with TNA. Then, we found the conditions to resolubilize EHEP–TNA precipitate for EHEP recycling. The obtained *aku*BGL structure suggests that only one GH1D (GH1D2) possesses laminarin hydrolytic activity; subsequently, ligand-docking experiments demonstrated that TNA/phlorotannin has a higher docking score than laminarin. Our results revealed the mechanisms by which EHEP protects *aku*BGL from phlorotannin inhibition and phlorotannin inhibits the hydrolytic activity of *aku*BGL, providing structural support for the potential application of brown algae for biofuel production.

## Results

### Effects of TNA on *aku*BGL activity with or without EHEP

Phlorotannin, a type of tannin, is a chemical defense metabolite of brown algae. It is difficult to isolate a compound from phlorotannins because they are a group of polyphenolic compounds with different sizes and varying numbers of phloroglucinol units ^13^, such as eckol, dieckol, and so on ^14^. Previous studies have reported that the phlorotannin-analog TNA has a comparable inhibition effect on *aku*BGL to that of phlorotannin ^4^. Hence, we used TNA instead of a phlorotannin to explore phlorotannin binding with EHEP and *aku*BGL. This activity assay system involves multiple equilibration processes: *aku*BGL⇋substrate, *aku*BGL⇋TNA, and EHEP ⇋TNA. First, we confirmed TNA inhibition of *aku*BGL activity and clarified the protective effects of EHEP from TNA inhibition. The inhibition experiments showed that the galactoside hydrolytic activity of *aku*BGL decreased with increasing TNA concentration, indicating that TNA inhibits *aku*BGL activity in a dose-dependent manner (Fig. 1A, Fig. S1A). Approximately 70% *aku*BGL activity was inhibited at a TNA concentration of 40 μM. Moreover, protection ability analysis revealed that EHEP protects *aku*BGL activity from TNA inhibition in a dose-dependent manner, as indicated by the recovery of the inhibited *aku*BGL activity with increasing EHEP concentration (Fig. 1B, Fig. S1B). Further, approximately 80% of *aku*BGL activity was recovered at an EHEP concentration of 3.36 μM.

**Figure 1.**
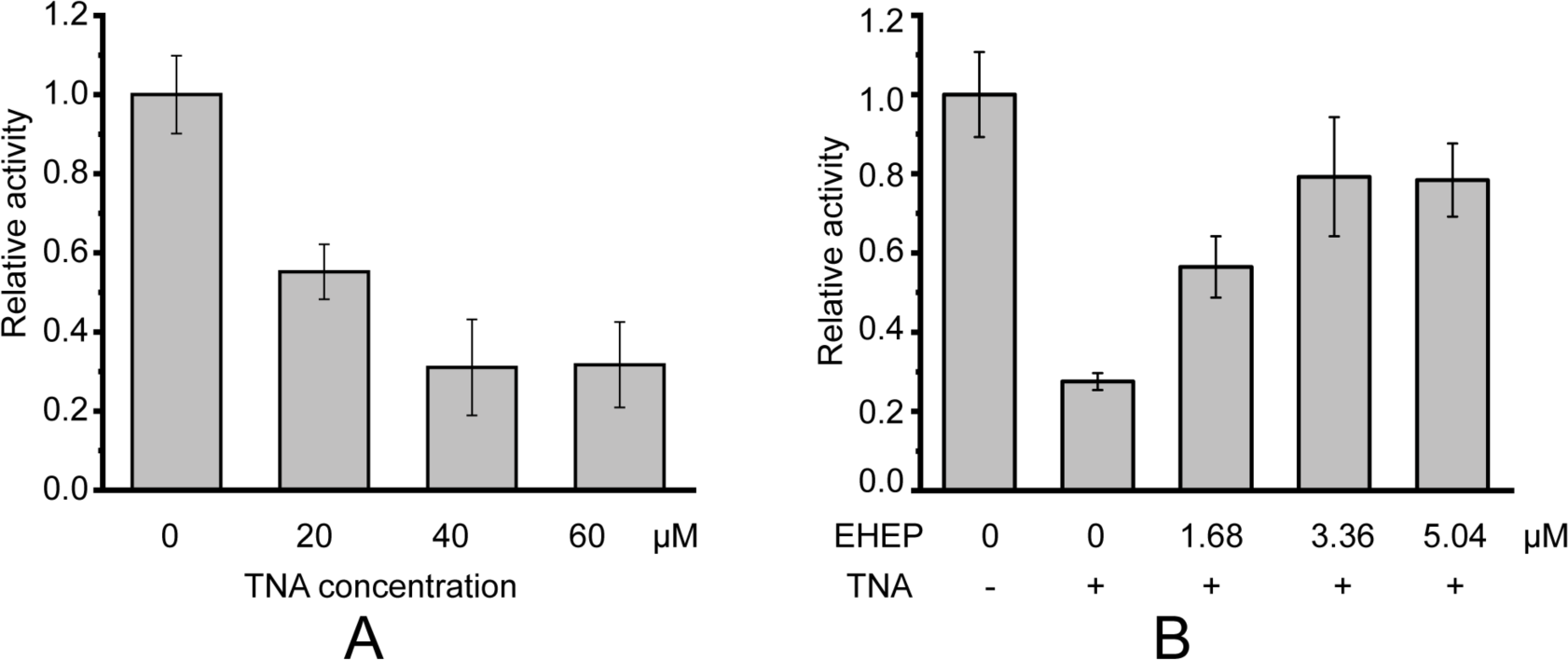
Galactoside hydrolytic activity of *aku*BGL toward ortho-Nitrophenyl-β-galactoside. Activity (%) is shown as the fold increase relative to *aku*BGL without the addition of TNA nor EHEP. (A) The hydrolytic activity of *aku*BGL (0.049 μM) with TNA at different concentrations. (B) The hydrolytic activity of *aku*BGL (0.049 μM) with 40 μM TNA and EHEP at different concentrations. The average and standard deviation of the relative activity were estimated from three independent replicates (N = 3).

### Overall structure of EHEP

Considering the lack of known homologous proteins of EHEP, we determined the structure of EHEP using the native-SAD method at a resolution of 1.15 Å, with a R_work_ and R_free_ of 0.18 and 0.19, respectively (Table 1). The residues A21–V227 in purified EHEP (1–20 aa were cleaved during maturation) were built, whereas two C-terminal residues were disordered. The structure of EHEP consists of three PADs: PAD1 (N24– C79), PAD2 (I92–C146), and PAD3 (F164–C221), which are linked by two long loops, LL1 (Q80–N91) and LL2 (R147– G163), and arranged in a triangle shape (Fig. 2A). These three PADs share a similar structure, with a root-mean-squared difference (RMSD) of 1.065 Å over 46 Cα atoms and only ∼20.3% sequence identity (Figs. 2B and 2C). The three PADs share a canonical CBM14 fold consisting of two β-sheets containing three N-terminal and two C-terminal antiparallel β-strands. Additionally, two small α-helices were appended to the N- and C-terminus in PAD1 and PAD3 but not in PAD2 (Fig. 2B).

**Figure 2.**
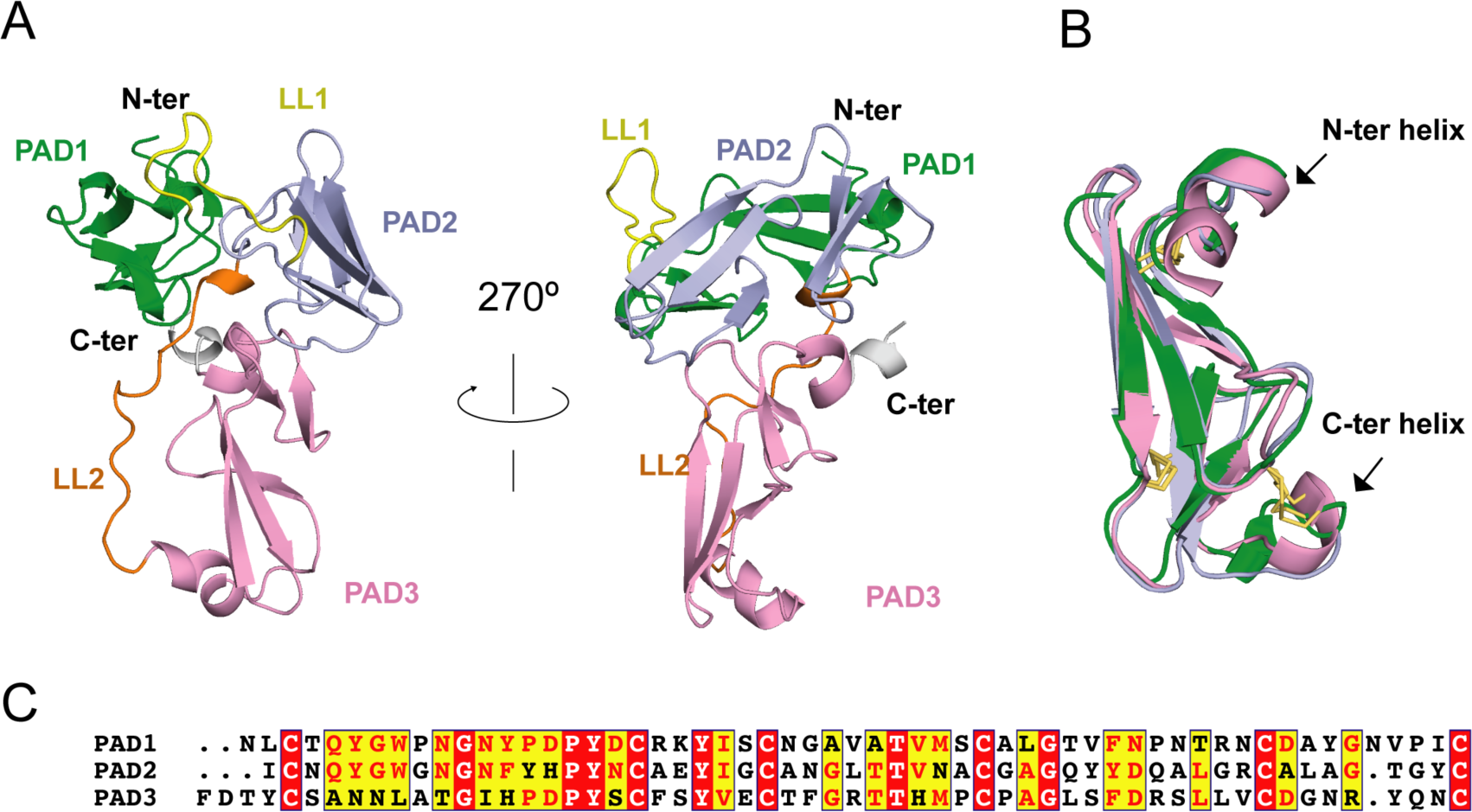
EHEP structure. (A) Cartoon representation of EHEP. The three PAD domains are colored green, light blue, and pink, respectively. Linker long loop1 (LL1) and loop2 (LL2) are colored yellow and blue. (B) Structural superposition of the three PAD domains of EHEP. The three domains are colored as in (A). The disulfide bonds are shown as yellow sticks. (C) Sequence alignment of three PAD domains. Alignment was performed by CLUSTALW and displayed with ESPript3.

**Table 1.**
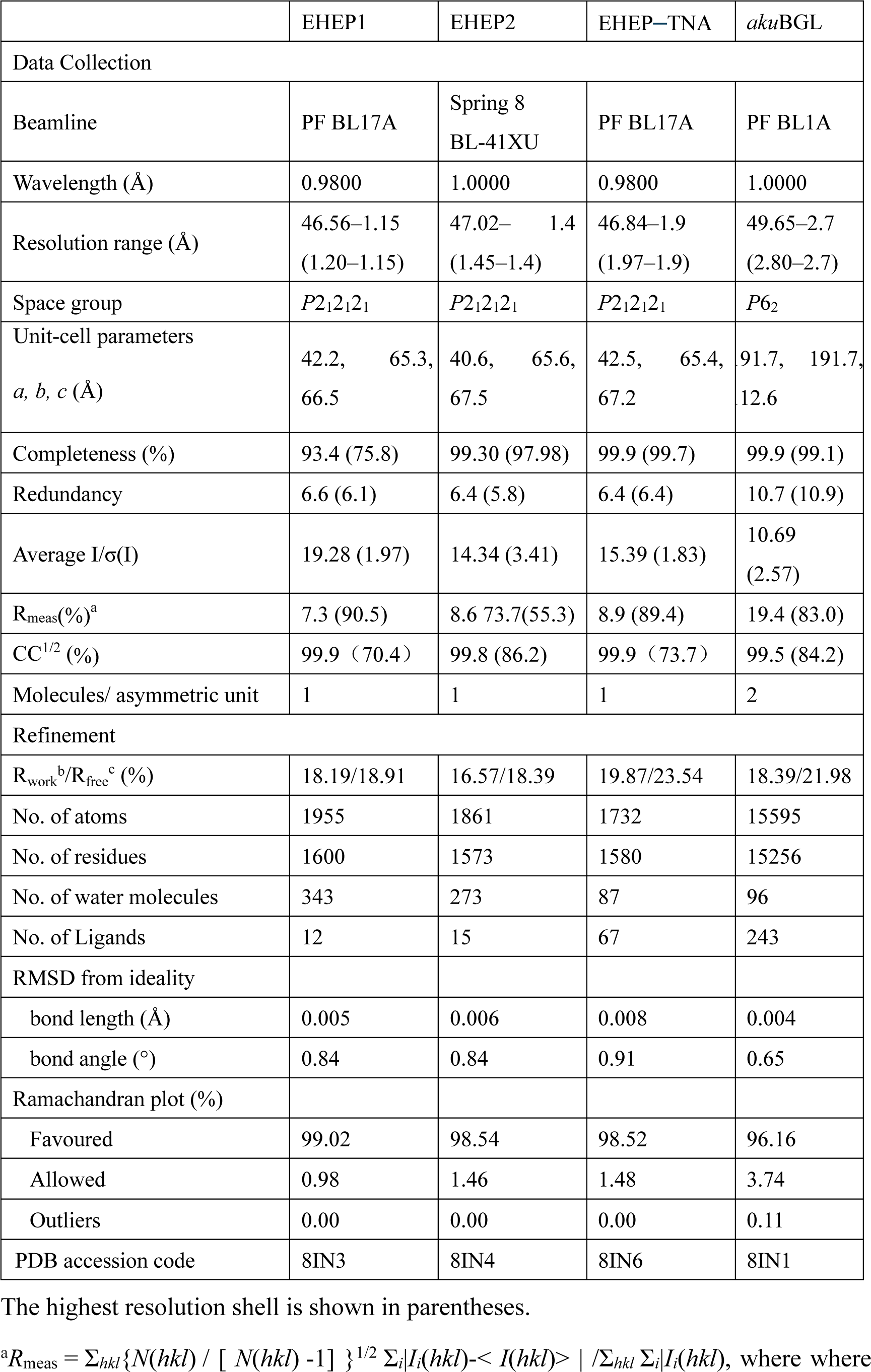
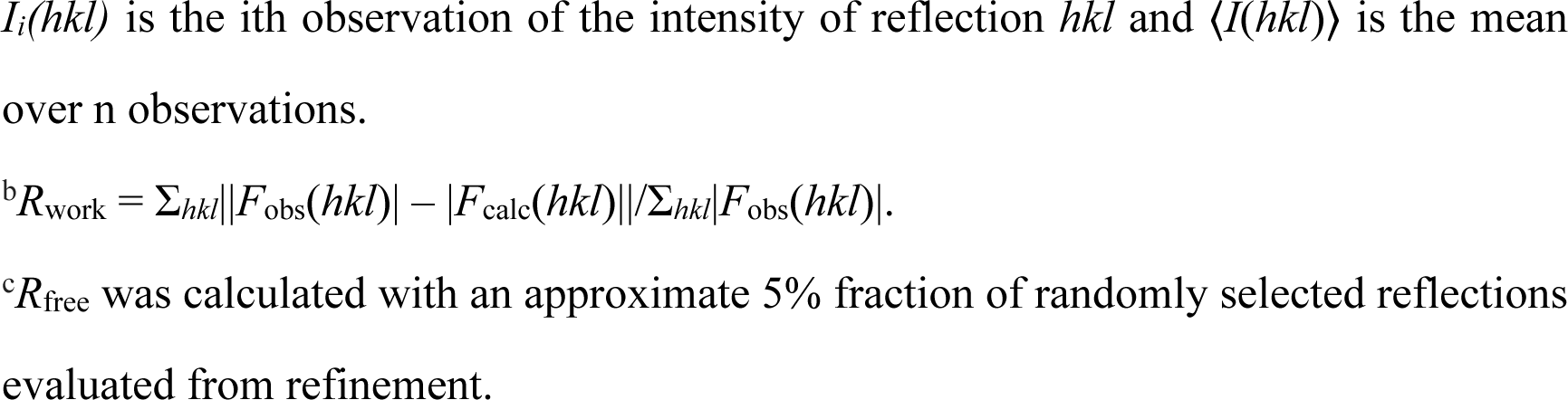
X-ray data collection and structure-refinement statistics.

|  | EHEP1 | EHEP2 | EHEP-TNA | akuBGL |
| --- | --- | --- | --- | --- |
| Data Collection |  |  |  |  |
| Beamline | PF BL17A | Spring 8<br>BL-41XU | PF BL17A | PF BL1A |
| Wavelength (Å) | 0.9800 | 1.0000 | 0.9800 | 1.0000 |
| Resolution range (Å) | 46.56–1.15<br>(1.20–1.15) | 47.02– 1.4<br>(1.45–1.4) | 46.84–1.9<br>(1.97–1.9) | 49.65–2.7<br>(2.80–2.7) |
| Space group | *P*2<sub>1</sub>2<sub>1</sub>2<sub>1</sub> | *P*2<sub>1</sub>2<sub>1</sub>2<sub>1</sub> | *P*2<sub>1</sub>2<sub>1</sub>2<sub>1</sub> | *P*6<sub>2</sub> |
| Unit-cell parameters<br>*a*, *b*, *c* (Å) | 42.2, 65.3,<br>66.5 | 40.6, 65.6,<br>67.5 | 42.5, 65.4,<br>67.2 | 91.7, 191.7,<br>12.6 |
| Completeness (%) | 93.4 (75.8) | 99.30 (97.98) | 99.9 (99.7) | 99.9 (99.1) |
| Redundancy | 6.6 (6.1) | 6.4 (5.8) | 6.4 (6.4) | 10.7 (10.9) |
| Average *I*/σ(*I*) | 19.28 (1.97) | 14.34 (3.41) | 15.39 (1.83) | 10.69<br>(2.57) |
| *R*<sub>meas</sub>(%)<sup>a</sup> | 7.3 (90.5) | 8.6 73.7(55.3) | 8.9 (89.4) | 19.4 (83.0) |
| CC<sup>1/2</sup> (%) | 99.9 (70.4) | 99.8 (86.2) | 99.9 (73.7) | 99.5 (84.2) |
| Molecules/ asymmetric unit | 1 | 1 | 1 | 2 |
| Refinement |  |  |  |  |
| *R*<sub>work</sub><sup>b</sup>/*R*<sub>free</sub><sup>c</sup> (%) | 18.19/18.91 | 16.57/18.39 | 19.87/23.54 | 18.39/21.98 |
| No. of atoms | 1955 | 1861 | 1732 | 15595 |
| No. of residues | 1600 | 1573 | 1580 | 15256 |
| No. of water molecules | 343 | 273 | 87 | 96 |
| No. of Ligands | 12 | 15 | 67 | 243 |
| RMSD from ideality |  |  |  |  |
| bond length (Å) | 0.005 | 0.006 | 0.008 | 0.004 |
| bond angle (°) | 0.84 | 0.84 | 0.91 | 0.65 |
| Ramachandran plot (%) |  |  |  |  |
| Favoured | 99.02 | 98.54 | 98.52 | 96.16 |
| Allowed | 0.98 | 1.46 | 1.48 | 3.74 |
| Outliers | 0.00 | 0.00 | 0.00 | 0.11 |
| PDB accession code | 8IN3 | 8IN4 | 8IN6 | 8IN1 |

The highest resolution shell is shown in parentheses.
<sup>a</sup>*R*<sub>meas</sub> = $\sum_{hkl} \{ N(hkl) / [ N(hkl) - 1 ] \}^{1/2} \sum_i |I_i(hkl) - \langle I(hkl) \rangle| / \sum_{hkl} \sum_i |I_i(hkl)|$ , where where
$I_i(hkl)$ is the $i$ th observation of the intensity of reflection $hkl$ and $\langle I(hkl) \rangle$ is the mean over $n$ observations.
${}^bR_{\text{work}} = \sum_{hkl} ||F_{\text{obs}}(hkl)| - |F_{\text{calc}}(hkl)|| / \sum_{hkl} |F_{\text{obs}}(hkl)|.$
${}^cR_{\text{free}}$ was calculated with an approximate 5% fraction of randomly selected reflections evaluated from refinement.

Although the Dali server ^15^ did not provide similar structures using the overall structure of EHEP as the search model, six structures showed similarities with a single PAD of EHEP. These structures were the members of the PAD family, including the chitin-binding domain of chitotriosidase (PDB ID 5HBF) ^16^, avirulence protein 4 from *Pseudocercospora fuligena* [*Pf*Avr4 (PDB ID 4Z4A)] ^17^ and *Cladosporium fulvum* [*Cf*Avr4 (PDBID 6BN0)] ^18^, allergen Der p 23 (PDB ID 4ZCE) ^19^, tachytitin (PDB ID 1DQC) ^20^, and allergen Blot 12 (PDB ID 2MFK), with Z-scores of 4.7–8.4, RMSD values of 1.2–2.8 Å, and sequence identity of 19%–37%. The highest sequence disparity was detected in PAD2, whereas the greatest structural differences were noted in PAD3. The C^No1^X_15_C^No2^X_5_C^No3^X_9_C^No4^X_12_C^No5^X_5-9_C^No^^6^ motif (superscripts and subscripts indicate the cysteine number and number of residues between adjacent cysteines, respectively) in each PAD of EHEP formed three disulfide bonds between the following pairs: C^No1^–C^No3^, C^No2^–C^No6^, and C^No4^–C^No5^ (Fig. 2B). Such rich disulfide bonds may play a folding role in the structural formation of EHEP, with >70% of the backbone in a loop conformation. A similar motif with disulfide bonds was observed in tachycitin ^20^, *Pf*Avr4 ^17^, *Cf*Avr4 ^18^, and the chitin-binding domain of chitinase ^16^. Although these proteins share a highly conserved core structure, they have different biochemical characteristics. For example, the chitin-binding domain of human chitotriosidase, Avr4 and tachycitin possess chitin-binding activity, but the critical residues for chitin-binding are not conserved ^16, 18, 21^, indicating that they employ different binding mechanisms. In contrast, EHEP and allergen Der p 23 do not possess chitin-binding activity ^4, 19^. Thus, the PAD family may participate in several biochemical functions.

### Modification of EHEP

Consistent with the molecular weight results obtained using MALDI–TOF MS ^22^, the apo structure2 (1.4 Å resolution) clearly showed that the cleaved N-terminus of Ala21 underwent acetylation (Fig. S2A), demonstrating that EHEP is acetylated in *A. kurodai* digestive fluid. N-terminal acetylation is a common modification in eukaryotic proteins. Such acetylation is associated with various biological functions, such as protein half-life regulation, protein secretion, protein–protein interaction, protein–lipid interaction ^23^, metabolism, and apoptosis ^24^. Further, N-terminal acetylation may stabilize proteins ^25^. To explore whether acetylation affects the protective effects of EHEP on *aku*BGL, we used the *E.coli* expression system to obtain the unmodified recomEHEP (A21–K229). We measured the TNA-precipitating assay of recomEHEP. The results revealed that recomEHEP precipitated after incubation with TNA, at a comparable level to that of natural EHEP (Fig. S2B), indicating that acetylation is not indispensable for the phlorotannin binding activity and stabilization of EHEP. Future studies are warranted to verify the exact role of N-terminal acetylation of EHEP in *A.kurodai*.

### TNA binding to EHEP

To understand the mechanism by which TNA binds to EHEP, we determined the structure of EHEP complexed with TNA (EHEP–TNA) using the soaking method. In the obtained structure, both 2F_o_–F_c_ and F_o_–F_c_ maps showed the electron density of 1,2,3,4,6-pentagalloylglucose, a core part of TNA missing the five external gallic acids (Fig. 3A, Fig. S3A). Previous studies have shown that acid catalytic hydrolysis of TNA requires a high temperature of 130°C ^26^; even with a polystyrene-hollow sphere catalyst, a temperature of 80°C is required ^27^. Therefore, the five gallic acids could not be visualized in the EHEP–TNA structure most likely due to the structural flexibility of TNA.

**Figure 3.**
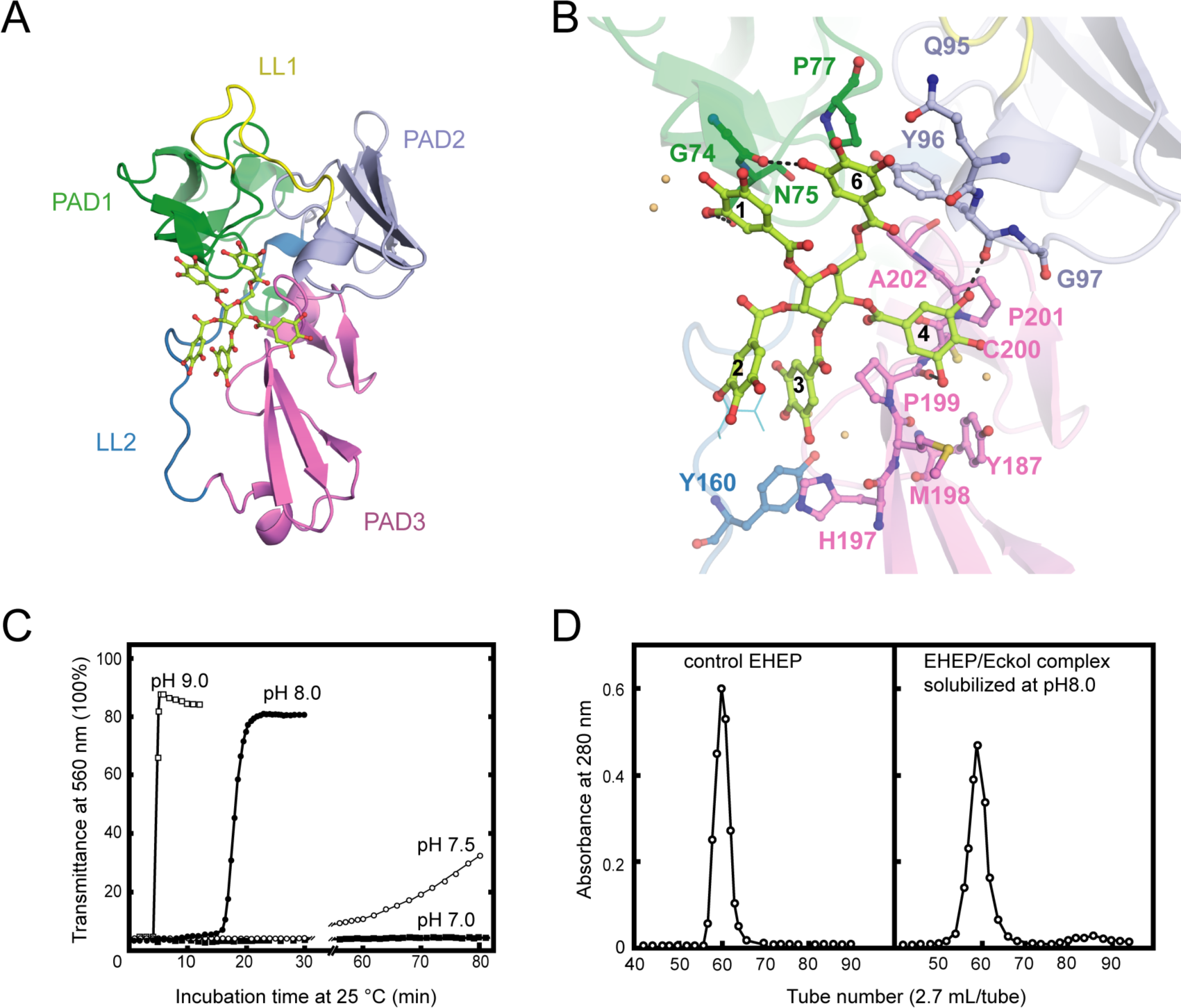
Structure of EHEP–TNA. (a) The overall structure of EHEP–TNA. (A) the overall structure of EHEP–TNA. EHEP and TNA are shown by the cartoon and stick model, respectively. EHEP is colored as in Fig. 1. The C and O atoms of TNA are colored lemon and red, respectively. (B) Interaction of TNA (ball-stick in same color as (A)) with EHEP (cartoon in same color as (A)) in EHEP–TNA structure. The residues of EHEP in contact are labeled and shown by a ball-stick with N, O, and S atoms in blue, red, and brown, respectively. The C and O atoms of TNA are colored the same as (A), lemon and red, respectively. Dashed lines show hydrogen bonds. The water molecules stabilizing TNA was shown as light orange spheres. (C) Effect of pH on resolubilization of an EHEP–eckol precipitate. Buffers with pH 9.0, 8.0, 7.5, and 7.0 are presented as hollow square, solid circle, hollow circle, and solid square, respectively. (D) The EHEP–eckol precipitate was dissolved in 50 mM Tris-HCl (pH 8.0) and analyzed using a gel filtration column of Sephacryl S-100.

The apo EHEP and EHEP–TNA structures were extremely similar, with an RMSD value of 0.283 Å for 207 Cα atoms (Fig. S3B). However, the superposition of the two structures showed a decrease of the loop part in EHEP–TNA. TNA binding caused a slight increase in the α-helix and β-sheet contents of PAD2 and PAD3 (Fig. S3B). In the EHEP–TNA structure, the residues C93–Y96 of PAD2 folded into an α-helix and each β-sheet of the first β-strand in PAD3 elongated by incorporating one residue in the first (G^176^) and second β-sheets (S^186^) and three residues in the third β-sheet (H^197^MP^199^). The EHEP–TNA structure revealed that TNA binds to the center of the triangle formed by the three PADs, a positively charged surface (Fig. 3A and Fig. S3C). The binding pocket on EHEP surface was formed by the C-terminal α-helix of PAD1, the N-terminal α-helix of PAD2, and the middle part (loop) of PAD3 assisted by two long linker loops (LL1 and LL2). TNA was primarily bound to EHEP via hydrogen bonds and hydrophobic interactions (Fig. 3B). Gallic acid1, 4, and 6 interacted with EHEP via hydrogen bonds and additional hydrophobic contacts, whereas gallic acid2 and 3 only interacted hydrophobically with EHEP. The 3-hydroxyl groups of gallic acid1 and 6 individually formed a hydrogen bond with the main chain of G74 and the side chain of N75 in PAD1. The backbone carbonyl of Y96 and P199 in PAD2 and PAD3, respectively, formed a hydrogen bond with the 3,5-hydroxyl groups of gallic acid4. Additionally, some hydrogen bonds were formed between TNA and water molecules. TNA binding was also stabilized by hydrophobic interactions between the benzene rings of gallic acid and EHEP. For instance, gallic acid4 and 6 showed alkyl–π interaction with P201 and P77, respectively; moreover, gallic acid3 and 4 formed amide–π stackings with P199.

The EHEP–TNA structure clearly showed that TNA binds to EHEP without covalent bonds and the binding does not induce significant structural changes; thus, we attempted to recover EHEP from EHEP–TNA precipitates by adjusting the pH. As hypothesized, re-solubilization of the EHEP–phlorotannin precipitate is pH-dependent (Fig. 3C). The EHEP–TNA precipitate did not resolubilize at pH 7.0; however, after incubating for >1 h at pH 7.5, the precipitate started resolubilizing. Most of the precipitate rapidly resolubilized at an alkaline pH (≥8.0) after incubation for 15 min. Further, the resolubilized EHEP had the same elution profile as that of the natural EHEP (Fig. 3D) in SEC, suggesting that resolubilized EHEP maintained the native structure and its phlorotannin-precipitate activity (Fig. S3D).

### Two domains of *aku*BGL

To reveal the structural basis of *aku*BGL recognition of laminarin and inhibition by TNA, we attempted to determine its structure. We soaked crystals in TNA as well as various substrate solutions but finally obtained the optimal resolution using crystal soaking in TNA. There was no electron density of TNA or something similar in the 2Fo–Fc and Fo–Fc map of the obtained structure; thus, we considered this structure as the apo form of *aku*BGL.

Two *aku*BGL molecules were observed in an asymmetric unit (MolA and MolB). These molecules lacked the N-terminal 25 residues (M1–D25), as confirmed using N-terminal sequencing analysis of purified natural *aku*BGL. This N-terminal fragment was predicted to be a signal peptide using the web server SignalP-5.0. The residues L26–P978 were constructed in both MolA and MolB with glycosylation, whereas the remaining C-terminal residues (A979–M994) could not be visualized as they were disordered. The electron density map of Fo–Fc revealed N-glycosylation at three residues, i.e., N113, N212, and N645 (Figs. S4A, B). N-glycosylation of GH enzymes prevents proteolysis and increases thermal stability ^28, 29^. Additionally, a study on β-glucosidase *Aspergillus terreus* BGL demonstrated that N-glycosylation of N224 affected the folding stability, even when it is located close to a catalytic residue ^30^. In *aku*BGL, all N-glycosylation sites were present on the surface, far away from the catalytic site. Therefore, we speculate that *aku*BGL glycosylation does not affect its activity. Except for the difference in visualized glycans resulting from glycosylation, MolA and MolB were similar, with a RMSD value of 0.182 for 899 Ca atoms; therefore, we used MolA for further descriptions and calculations.

The structure of *aku*BGL consisted of two independent GH1 domains, GH1D1 (L26–T494) and GH1D2 (D513–P978), linked by a long loop (D495–Y512) (Fig. 4A). There was little interaction between GH1D1 and GH1D2, only in a buried surface area comprising 2% of the total surface (708.9 Å^2^) (Fig. S4C). GH1D1 and GH1D2 have a sequence identity of 40.47% and high structural similarity with an RMSD value of 0.59 Å for 371 Cα atoms (Fig. S5A upper panel).

**Figure 4.**
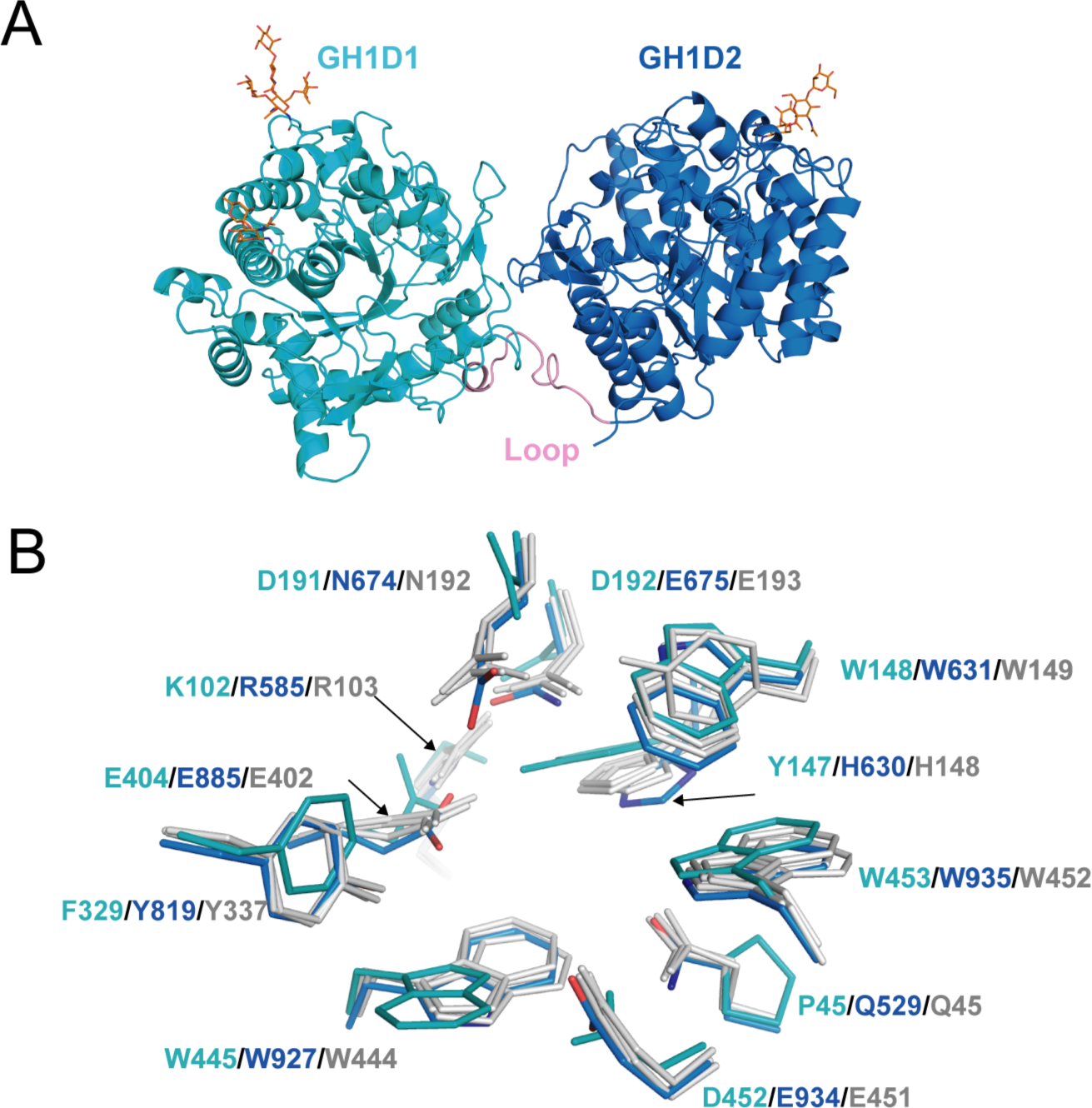
Structure of *aku*BGL. (A) Overall structure. The GH1D1 (light blue) and GH1D2 (cyan) domains are linked by a long loop (linker-loop) colored in pink. The N-linked glycans were shown in the orange stick. (B) Residues superposition of the GBS and CR sites of the domains GH1D1 (cyan), GH1D2 (light blue) with β-glucosidase structures from termite *Neotermes koshunensis* (*Nk*BGL, grey(PDB ID 3VIH) ^32^, β-glucosidase from rice (*Os*BGL, gray, PDB ID 2RGL) ^35^, β-glucosidase from *Bacillus circulans* sp. *Alkalophilus* (grey, PDB ID 1QOX) ^67^. Only the residues numbers of GH1D1 (cyan), GH1D2 (light blue), and *Nk*BGL (grey) are shown for clarity.

Glucosidases of the GH1 family utilize the retaining mechanism with two glutamic acids for catalyzing glucoside hydrolysis. In general, the distance between the two catalytic oxygen atoms of the side chain of two glutamic acids is approximately 5 Å ^31^. Sequence and structure alignment of GH1D1 and GH1D2 of *aku*BGL with other members of the GH1 family revealed that the second glutamate is conserved (E404), but the first glutamate is replaced by D192 in GH1D1. The oxygen atoms of the side chains between D192 and E404 of GHD1 were 8.4 Å apart. In contrast, GH1D2 conserved two glutamic acids (E675 and E885) at the carboxyl termini of β-strands 4 and 7; the distance between oxygen atoms of E675 and E885 side chains was 5.1 Å (Fig. S5A bottom panel), similar to that of *Neotermes koshunensis* BGL (*Nk*BGL, PDB ID 3VIH) ^32^, *Nannochloropsis oceanica* BGL (*No*BGL, PDB ID 5YJ7) ^33^, and *Spodoptera frugiperda* BGL (PDB ID 5CG0) (3.9–4.9Å) ^34^. Furthermore, regarding the two other conserved essential regions for β-glucosidase activity, namely, glycone-binding site (GBS) and catalysis-related residues (CR), GH1D1 conserved neither GBS nor CR, whereas GH1D2 conserved both (Fig. 4B). Altogether, we suggest that GH1D1 does not possess catalytic activity. We expressed and purified the recombinant GH1D1, which did not show any hydrolytic activity toward O-PNG (Figs. S4D, E), although we could not rule out the effect of N-glycosylation.

Structural comparison of GH1D2 with other BGLs, including *Nk*BGL (PDB ID 3VIH) ^32^, rice (*Oryza sativa L.)* BGL (*Os*BGL, PDB ID 2RGL) ^35^, and microalgae *No*BGL (PDB ID 5YJ7) ^33^, revealed the characteristics of each active pocket (Fig. S5B). *Os*BGL and *No*BGL have a deep, narrow, and straight pocket, whereas GH1D2 and *Nk*BGL have a broad and crooked pocket. Such active pocket shapes reflect the substrate preferences of *Os*BGL and *No*BGL; they hydrolyze laminaribiose with no detectable activity toward laminaritetraose ^33, 36^. Furthermore, the difference in the features of large active pockets between *Nk*BGL and GH1D2, wherein GH1D2 often possesses an auxiliary site with several aromatic residues bound to the carbohydrate via CH–π interactions ^37^, may explain their substrate specificity. *Nk*BGL efficiently hydrolyzes laminaribiose and cellobiose but has weak hydrolytic activity toward laminarin ^38^. In contrast, the GH1D2 of *aku*BGL has similar activity levels toward cellobiose and laminarin ^39^. Therefore, the GH1D2 of *aku*BGL may recognize larger substrates than that of other BGLs. Laminarin typically has a curved conformation; accordingly, narrow- and straight-shaped pockets are incompatible for binding. Furthermore, we docked GH1D2 with laminaritetraose, wherein the four glucose units formed extensive contacts with GH1D2. Hydrogen bonds involved the catalytic residues E675. In addition, several aromatic residues, such as W631, F677, W681, F689, Y819, Y846, W857, and W935, formed CH–π stacking (Fig. S6). Some interacting residues belonged to GBS and CR sites, such as E675, W631, Y819, and W935. Additionally, the docking structure revealed that the +3 and +4 glucose of laminaritetraose is located at the auxiliary binding site and that atom O1 of the +4 glucose is positioned outside the pocket (Fig. S6), implying that the auxiliary binding site with several aromatic residues (F677 and W681) of GH1D2 facilitates laminarin binding.

### Inhibitor binding of *aku*BGL

As we could not obtain the complex structure of *aku*BGL with TNA, we performed docking calculations of *aku*BGL GH1D2 with TNA to explore the inhibition mechanism. The docking model of *aku*BGL–TNA showed that seven gallic acid rings of TNA formed an extensive hydrogen bond network with *aku*BGL in the binding pocket (Fig. 5). The hydroxyl groups of TNA formed hydrogen bonds with the residues N552, E675, D735, K739, K759, Q840, T844, D852, and K859 of GH1D2. Moreover, benzene rings showed hydrophobic interactions with several hydrophobic residues. In particular, stable π–π stacking was observed between TNA and residues F547, W631, F689, Y846, W857, and W935. Among these residues, the conserved E675 was the catalytic residue, and W631, W935, and E934 contributed to GBS and CR sites.

**Figure 5.**
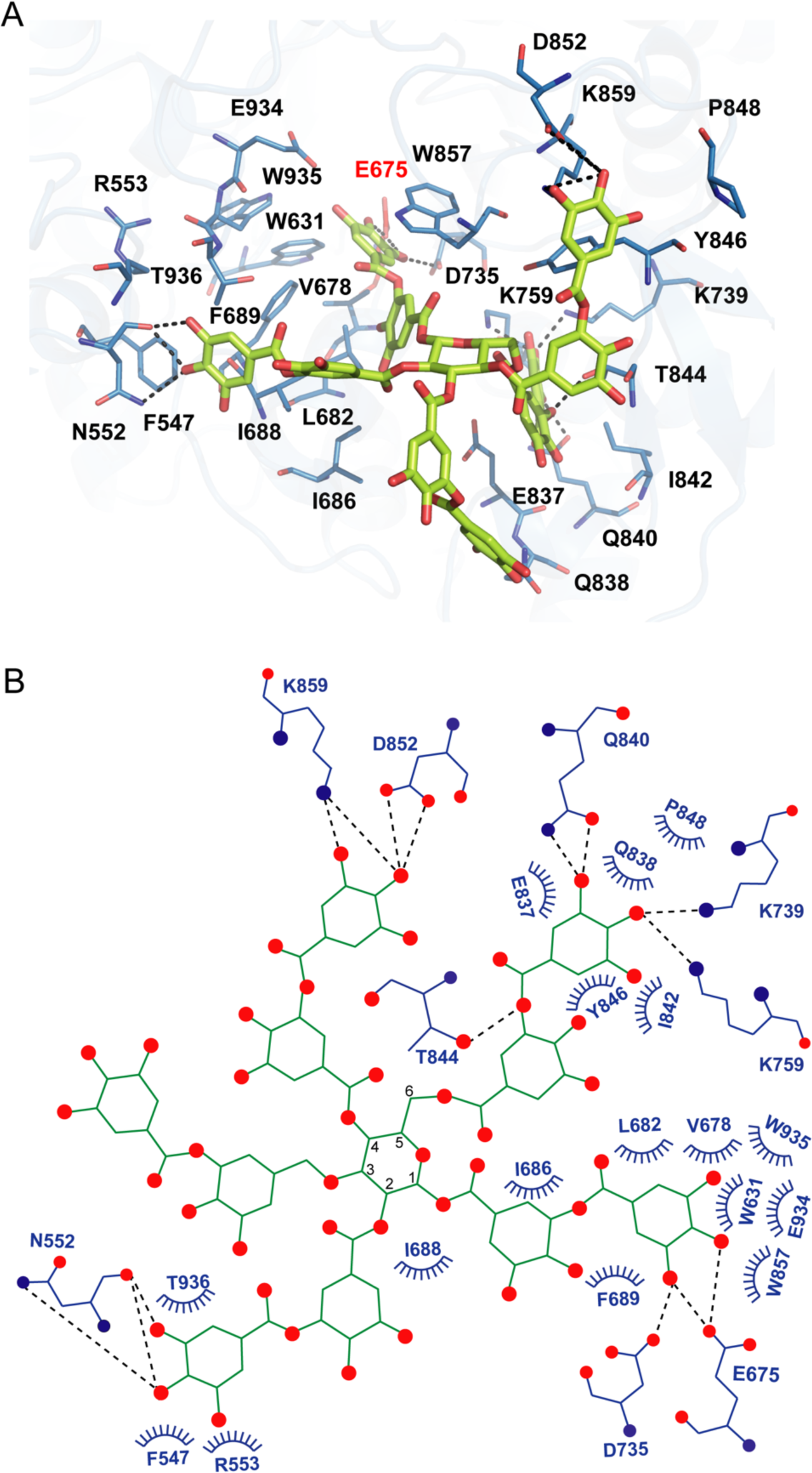
Docking model of *aku*BGL with TNA. (A) Detailed interaction between *aku*BGL and TNA in the docking model. TNA is shown in the green stick model. The hydrogen bonds are shown as dashed lines. (B) A 2D diagram of the interaction between *aku*BGL and TNA shown in (A). The hydrogen bonds are shown as dash lines, and the hydrophobic contacts as circular arcs.

In addition, we performed a docking calculation of GH1D2 with the characteristic inhibitors eckol and phloroglucinol ^40^. The binding mechanisms of eckol and phloroglucinol were similar to those of TNA but with different contact residues (Fig. S7). For eckol, the six hydroxyl groups formed hydrogen bonds with residues E675, D735, E737, K759, E885, and E934. Additionally, residues W631, F677, F689, Y819, W857, W935, F943, and W927 formed π–π stacking with eckol. For phloroglucinol, the three hydroxyl groups formed hydrogen bonds with E675, E885, and E934, whereas residues W631, F689, Y819, W857, W935, and W927 formed π–π stacking with the benzene ring.

The docking scores of the inhibitors TNA, eckol, and phloroglucinol were −8.8, −7.3, and −5.7 kcal/mol, respectively, whereas the substrate laminaritetraose had a docking score of −6.6 kcal/mol. Those docking scores corroborated well with the inhibition activity toward *aku*BGL, that TNA had a more robust inhibition activity than phloroglucinol ^4^, indicating that the docking results are reasonable. In summary, the three inhibitors interacted with *aku*BGL through similar binding mechanisms to occupy the substrate-binding site, suggesting a reversible competitive inhibition mechanism.

## Discussion

In marine habitats, the ecological interactions between brown algae and herbivores dominate marine ecosystems ^41^. The *aku*BGL–phlorotannin/laminarin–EHEP system represents the feeding defense–offense associations between *A. kurodai* and brown algae. We focused on this system to understand the molecular mechanism at the atomic level. In contrast to most GH1 BGLs containing one catalytic GH1 domain, *aku*BGL consists of noncatalytic GH1D1 and catalytic GH1D2. The noncatalytic GH1D1 may act as a chaperone of GH1D2, as we successfully overexpressed GH1D1 but failed to do the same for GH1D2. Such multi-GH1D assembly and a similar function has been suggested in β-glucosidase *Cj*CEL1A of *Corbicula japonica* ^42^ and glycosidase *LpM*DGH1 of the shipworm *Lyrodus pedicellatus*^43^. *Cj*CEL1A has two tandem GH1Ds with a sequence identity of 43.41% ^42^. Two catalytic glutamic acids and the residues related to substrate binding are conserved in the second GH1D, whereas the first GH1 domain lacks these conserved residues and may play a role in folding the catalytic domain. *LpM*DGH1 consists of six GH1Ds, among which GH1D2, 4, 5, and 6 contain the conserved residues for activity, whereas others do not contain these residues and might be involved in protein folding or substrate interactions ^43^. Future assay of GH1D2 and inactive mutants is the complement to validate the molecular mechanism of *aku*BGL.

BGLs have different substrate preferences in the degree of polymerization and type of glycosidic bond. In general, BGLs prefer to react with mono-oligo sugars over polysaccharides. For instance, *Os*BGL, *No*BGL, and *Nk*BGL hydrolyze disaccharides (cellobiose and laminaribiose) but display no or weak activity toward polysaccharides (cellulose and laminarin) ^33, 36, 38^. The structure of GH1D2 explained the substrate preference for the polysaccharide laminarin. GH1D2 contains an additional auxiliary site composed of aromatic residues (Fig. S5B) in the substrate entrance pocket, which putatively enables it to accommodate a long substrate, contributing to *aku*BGL activity toward laminarin, as supported by docking calculations (Fig. S6). In addition, docking analysis of *aku*BGL GH1D2 with inhibitors (TNA, phlorotannin, eckol, and phloroglucinol) revealed that these inhibitors bind to the substrate-binding site via hydrogen bonds and hydrophobic interactions similar to laminarin. Such binding mechanisms suggest the presence of competitive inhibition to occupy the binding site, consistent with previous research ^4^. Future kinetic experiments are required to validate quantitatively the competitive inhibition of phlorotannin against *aku*BGL.

EHEP, expressed in the midgut of *A. kurodai*, was identified as an antidefense protein, protecting the hydrolysis activity of *aku*BGL from phlorotannin inhibition ^4^. Such an ecological balance also exists between plants and their predatory mammals and insects. Similar to brown algae, plants use the toxic secondary metabolite tannins as their defense mechanism against predators, which constitute 5%–10% of dry weight of leaves. In vertebrate herbivores, tannins reduce protein digestion ^44^. In phytophagous insects, tannins may be oxidized at the alkaline pH of the insect midgut and cause damage to cells ^45^. The evolution of plant–herbivore survival competition has led to the development of remarkably unique adaptation strategies. Mammals feeding on plants that contain tannin may overcome this defense by producing tannin-binding proteins, proline-rich proteins, and histatins ^46 47^. Proline constitutes at least 20% of the total amino acid content in proline-rich proteins; for some species, the proportion of proline reaches 40%. Histidine constitutes 25% of total amino acid content in histatins. Both proline-rich proteins and histatins are unfolding proteins with randomly coils in solution. In caterpillars, the oxidation damage of tannin is reduced by the low oxygen level. Some insects use the peritrophic membrane to transport tannins into the hemolymph, where they are excreted ^48^. Additionally, the peritrophic envelop protects insects from tannins by forming an impermeable barrier to tannins ^44^. *A. kurodai* uses a similar strategy to mammals by secreting the tannin-binding protein EHEP. Although EHEP has a completely different amino acid composition with proline-rich proteins and histatins, EHEP also binds to phlorotannin. Therefore, EHEP may be a specific counteradaptation that allows *A.kurodai* to feed on brown algae, as there are no homologous proteins in other organisms.

The three PADs of EHEP are arranged in a triangle shape, forming a large cavity on the surface at the triangle center to provide a ligand-binding site. EHEP has a positively charged surface at a pH of <6.0, whereas the surface becomes negatively charged at a pH of >7.0 (Fig. S3C). Meanwhile, TNA has a pKa of 4.9–8 ^49–51^, showing minor negative charges at an acidic pH and the highest negative charge at a pH of >7.0 ^52^. Therefore, TNA binds to EHEP at a pH of <6.0 (pH of crystallization = 4.5), but it shows charge repulsion with EHEP at a pH of >8.0. Altogether, TNA is protonated and behaves as a hydrogen bond donor when the pH is below its pKa, whereas when the pH is above its pKa, TNA is deprotonated and the hydrogen bonding cannot be maintained. As losing hydrogen bonds and increasing repulsive forces at a pH >8.0, the precipitated EHEP–TNA could dissolve in the buffer of pH >8.0. This pH-induced reversible interaction also occurred in other proteins, such as BSA, pepsin, and cytochrome C ^53^. The phlorotannin members share a similar structure with TNA; thus, we speculate that the EHEP–phlorotannin complex also exhibits a pH-induced reversible interaction. *In vivo*, the pH of the digestive fluid of *A. kurodai* is approximately 5.5^4^, which favors the binding of EHEP to phlorotannin. In the alkaline hindgut ^54^, the EHEP–phlorotannin disassociates (Fig. 6), and the phlorotannin is subsequently excreted from the anus.

**Figure 6.**
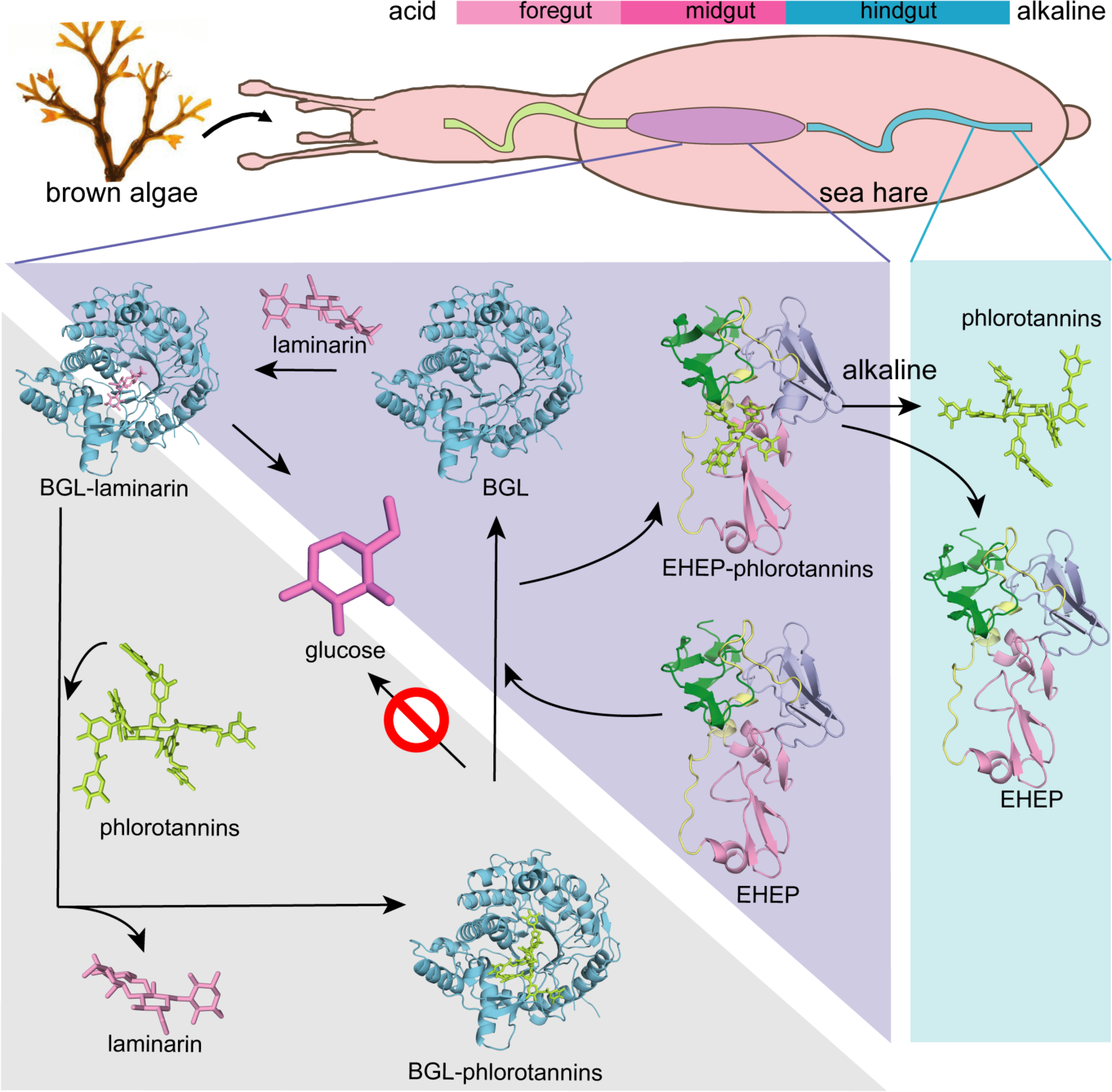
Proposed molecular mechanisms. Proposed molecular mechanisms of TNA inhibition of *aku*BGL activity and EHEP’s protective effects of *aku*BGL in *aku*BGL– phlorotannin/laminarin–EHEP system (light purple triangle). The digestive tract of *A.kurodai* consists of foregut (blue), midgut (purple), and hindgut (blue) (top). The bar chart above depicts the pH of the digestive tract, with pink denoting acid and blue denoting alkalinity.

Based on the EHEP–TNA structure and docking models of *aku*BGL– inhibitor/substrate, we proposed a mechanism of phlorotannin inhibition on *aku*BGL activity and EHEP protection from phlorotannin inhibition (Fig. 6). Because laminarin lacks the benzene rings essentially to form CH–π stacking interactions with EHEP, the EHEP can be considered not binding with the laminarin. In the absence of EHEP, phlorotannin occupies the substrate-binding site of *aku*BGL, inhibiting the substrate from entering the active site and resulting in no glucose production. In the presence of EHEP, it competitively binds to phlorotannin, freeing the *aku*BGL pocket. Then, the substrate can enter the active pocket of *aku*BGL and glucose can be produced normally. The digestive fluid of *A. kurodai* contains EHEP at a high concentration (>4.4 µM) ^4^, which is slightly higher than the concentration of EHEP (3.36 µM) that protects *aku*BGL activity (Fig. 1B). The high concentration of EHEP allows *A. kurodai* feeding of phlorotannin-rich brown algae. The balance between phlorotannin inhibition and protection is controlled by the concentrations of phlorotannin and EHEP *in vivo*.

The *aku*BGL–phlorotannin/laminarin–EHEP system is the digestive–defensive– offensive associations between algae and herbivores. Our study presented the molecular mechanism of this system at the atomic level, providing a molecular explanation for how the sea hare *A. kurodai* utilizes EHEP to protect *aku*BGL activity from phlorotannin inhibition. Further, such a feeding strategy has attracted attention for producing glucose as a renewable biofuel source, so our studies provide a molecular basis for the biofuel industry applications of brown algae.

## Materials and Methods

### EHEP and *aku*BGL preparation

Natural EHEP (22.5 kDa) and *aku*BGL (110 kDa) were purified from *A. kurodai* digestive fluid as described previously ^4^. For crystallization, we further added one step for purification of EHEP using size exclusion chromatography (HiLoad 16/60 Superdex 75, GE Healthcare), for which the column was equilibrated with 20 mM MES-NaOH buffer (pH 6.5). Obtained EHEP was then concentrated to 15–25 mg/mL using Vivaspin-4 10K columns (Sartorius, Göttingen, Germany). About *aku*BGL, we exchanged buffer from 20 mM Tris-HCl pH 7.0 to 20 mM Bis-tris pH 6.0 and concentrated it to 11 mg/mL using Amicon with a cutoff of 50 kDa.

To verify whether the chemical modification which was indicated by previous study^22^ affects the function of EHEP, we prepared recombinant EHEP (recomEHEP) without the N-terminal signal peptide (1–20 aa) and chemical modifications ^22^. EHEP cDNA was obtained via reverse transcription–polymerase chain reaction (RT–PCR) using the total RNA of *A. kurodai* as a template. The reamplified fragment was digested and ligated to a plasmid derived from pET28a (Novagene, Darmstadt, Germany). We transformed the plasmid containing recomEHEP into *E. coli* B834(DE3) pARE2 and expressed it with a C-terminal hexahistidine-tag. The cells were cultured in lysogeny broth (LB) medium with the antibiotics kanamycin (25 mg/L) and chloramphenicol (34 mg/L) until the optical density at 600 nm (OD_600_) reached 0.6. Subsequently, overexpression was induced by adding 0.5 mM isopropyl b-D-L-thiogalactopyranoside for 20 h at 20 ℃. After harvesting by centrifugation, the cells were resuspended in a buffer containing 50 mM Tris-HCl pH 7.4, 300 mM NaCl, DNase, and lysozyme and were disrupted via sonication. The insoluble part was removed by centrifugation for 30 min at 40000 ×*g* at 4 ℃. We loaded the supernatant onto a 5-mL Histrap HP column and the recomEHEP was eluted using increasing concentrations of imidazole (0–500 mM). The purified proteins were dialyzed against a solution containing 50 mM Tris-HCl pH 7.4 and 50 mM NaCl and subsequently loaded onto a Hitrap Q column and eluted by a linear gradient of a solution containing 50 mM Tris-HCl and 1 M NaCl. Fractions containing recomEHEP were concentrated and then purified using a gel filtration column (Hiload 16/60 superdex 75 pg) equilibrated with 20 mM sodium acetate pH 6.0 and 100 mM NaCl. We collected the fractions containing recomEHEP and concentrated them to 2.1 mg/mL using Amicon (Merck, American).

DNA encoding *aku*BGL without signal peptide was codon optimized (GENEWIZ, China) for overexpress in *E. coli*, and cDNA encoding GH1D1 domain was subcloned to a modified pET-32a vector (Invitrogen, USA) with an N-terminal TrxA tag and a 6×His tag followed by a TEV protease recognition site. The plasmid pET-32a-GH1D1 was electorally transformed into *E.coli* origami2 cells. We grew the cells in LB medium supplemented with 100 µg/mL of ampicillin at 37 °C until the OD_600_ reached 0.6. Then the cultures were cooled and induced with 0.5 mM isopropyl-β-D-thiogalactopyranoside at 20 ℃ for 16 h. After resuspending the harvested cells in a buffer containing 50 mM Tris-HCl (pH 7.4), 250 mM NaCl, and 5% glycerol, we disrupted the cells by sonication. The cell lysate was centrifugated at 40,000 ×g for 30 min at 4 ℃. The supernatant was filtered by 0.45 μm membrane and then loaded onto a Histrap HP column (GE, USA). After washing with lysis buffer supplemented with 20 mM imidazole, the recomGH1D1 was eluted linearly from the column with lysis buffer supplemented with 500 mM imidazole. The fractions containing recomGH1D1 were added with TEV protease at a ratio of 1:10 and dialyzed against the buffer containing 50 mM Tris (pH 7.4), 50 mM NaCl, and 2 mM DTT at 4 ℃ overnight. Then, we purified the recomGH1D1 by a Histrap HP column again, and collected the flowthrough. Finally, we purified the recomGH1D1 by size exclusion chromatography on a Hiload 16/60 200 pg column (GE, USA) equilibrated with a buffer containing 20 mM Bis-Tris (pH 6.0). The recomGH1D1 was concentrated to 1.5 mg/mL by centrifugation using Amicon Ultra with 10 kDa. cutoff.

### N-terminal sequencing of *aku*BGL

We performed a N-terminal sequencing of purified *aku*BGL using the Edman degradation method. *aku*BGL was separated by SDS–PAGE, followed by electrophoretic transfer onto a PVDF (polyvinylidene fluoride) membrane (GE, American). The membrane was subsequently stained with Ponceau S solution. The band corresponding to *aku*BGL was excised and analyzed using a PPSQ-53A Protein sequencer (Shimadzu, Japan) at the Instrumental Analysis Service of Hokkaido University.

### Effects of TNA on *aku*BGL activity with or without EHEP

Due to the yellow color of TNA, which affects absorbance at 420 nm of the reaction product o-nitrophenol of *aku*BGL, we used HPLC to measure the *aku*BGL activity in the reaction system involving TNA. Ortho-nitrophenyl-β-galactoside (ONPG) was used as a substrate to measure *aku*BGL activity. The reaction system (100 μL) included 2.5 mM ONPG, 49 nM *aku*BGL, and different TNA concentrations in a reaction buffer (50 mM CH_3_COONa pH 5.5, 100 mM NaCl, and 10 mM CaCl_2_). After incubation for 10 min at 37 ℃, 100 μL of methanol was added to each sample to terminate the reaction. Then, the mixture was centrifuged for 10 min at 15000 ×*g* at 4 ℃ and the supernatant was used for analyzing *aku*BGL activity via HPLC. To measure the protective effect of EHEP on *aku*BGL, we added different amounts of EHEP to the reaction system.

### RecomGH1D1 activity assay

We measured simply the activity of the recomGH1D1 using a spectrophotometer because the reaction product o-nitrophenol is yellow. The reaction system (100 μL) included 2.5 mM ONPG, 49 nM *aku*BGL or recomGH1D1 in a reaction buffer (50 mM CH_3_COONa pH 5.5, 100 mM NaCl, and 10 mM CaCl_2_). After incubation for 10 min at 37 ℃, the reaction was terminated by adding 100 μL of 500 mM Na_2_CO_3_. The absorbance at 420 nm was measured using a SpectraMax spectrophotometer (Molecular Devices, Japan).

### Binding assay for recomEHEP with TNA

We measured the binding activity of recomEHEP using precipitation analysis in same method of natural EHEP, as described previously ^4^. Briefly, recomEHEP or EHEP was incubated with TNA at 25 ℃ for 90 min and centrifuged for 10 min at 12000 ×*g* at 4 ℃. Then, we washed the precipitates twice and resuspended them in an SDS–PAGE loading buffer for binding analysis.

### Resolubilization of the EHEP–eckol precipitate

A mixture of 2 mg of EHEP and 0.4 mg of eckol was incubated at 37 °C for 1 h, followed by centrifugation at 12000 ×g for 10 min, and the supernatant was removed. The sediment was dissolved in 50 mM Tris-HCl buffer at different pH (7.0–9.0) and the absorbance at 560 nm was measured over time. After checking the elution peak by SDS–PAGE, the resolubilized EHEP was analyzed by a Sephacryl S-100 HR column (2.0 ×110 cm). Moreover, the eckol/dieckol-binding activity of resolubilized EHEP was assessed, as mentioned above in this section.

### Crystallization and data collection

The crystallization, data collection, and initial phase determination of EHEP were described previously ^22^. As EHEP precipitates when bound to TNA, we could not cocrystallize EHEP with TNA. Therefore, we used the soaking method to obtain the EHEP–TNA complex. Owing to the poor reproducibility of EHEP crystallization, we used a co-cage-1 nucleant ^55^ to prepare EHEP crystals for forming the complex with TNA. Finally, we obtained high-quality EHEP crystals under the reservoir solution containing 1.0 M sodium acetate, 0.1 M imidazole (pH 6.5) with co-cage-1 nucleant. Subsequently, we soaked the EHEP crystals in a reservoir solution containing 10 mM TNA at 37 ℃ for 2 days; then, they were maintained at 20 ℃ for 2 weeks. Next, we soaked the EHEP crystals in a reservoir solution containing 10 mM phloroglucinol. For data collection, the crystal was soaked in a cryoprotectant solution containing 20% (v/v) glycerol along with the reservoir solution. Diffraction data were collected under a cold nitrogen gas stream at 100 K using Photon Factory BL-17 (Tsukuba, Japan) or Spring 8 BL-41XU (Hyogo, Japan).

For *aku*BGL crystallization, the initial crystallization screening was performed using the sitting-drop vapor-diffusion method with Screen classics and Classics II crystallization kits (Qiagen, Hilden, Germany) and PACT kits (Molecular Dimensions, Anatrace, Inc.) at 20 ℃. Crystallization drops were set up by mixing 0.5 μL of protein solution with an equal volume of the reservoir solution. The initial crystals were obtained under condition no. 41 (0.1 M sodium acetate pH 4.5 and 25% polyethylene glycol [PEG] 3350) of Classics II, no. 13 (0.1 M MIB buffer [25 mM sodium malonate dibasic monohydrate, 37.5 mM imidazole, and 37.5 mM boric acid], with pH 4.0 and 25% PEG 1500), and no. 37 (0.1 M MMT buffer [20 mM DL-malic acid, 40 mM MES monohydrate, and 40 mM Tris], with pH 4.0 and 25% PEG 1500) of PACT. After optimization by varying the buffer pH and precipitant concentration and adding co-cage1 nucleant, the optimal crystals were obtained using 0.1 M sodium acetate pH 4.5, and 20% PEG 3350 with a co-cage1 nucleant at a protein concentration of 5.4 mg/mL. Diffraction data were collected under a cold nitrogen gas stream at 100 K using Photon Factory BL-1A (Tsukuba, Japan) after cryoprotection by adding glycerol to a 20% final concentration into the reservoir solution. The optimal resolution of diffraction data was obtained by soaking a crystal with 5 mM TNA in the reservoir buffer at 37 ℃ for 4 h.

All datasets were indexed, integrated, scaled, and merged using *XDS/XSCALE* program ^56^. Statistical data collection and process are summarized in Table 1.

### Structure determination and refinement

For EHEP structure determination, after initial phasing via the native-SAD method^22, 57^, the model was obtained and refined with *auto-building* using *Phenix.autobuild* of *Phenix* software suite ^58^. The obtained native-SAD structure was used as a model for rigid body refinement using *phenix.refine* ^59^ of *Phenix* software suite with a native data at high resolution of 1.15 Å. The structure of EHEP was automatically rebuilt using *Phenix.autobuild* of the *Phenix* software suite again ^58^. Several rounds of refinement were performed using *Phenix.refine* of the *Phenix* software suite ^58^, alternating with manual fitting and rebuilding using *COOT* program ^60^. The final refinement statistics and geometry are shown in Table 1.

The structure of the EHEP–TNA complex was determined via the molecular replacement (MR) method using the EHEP structure as a search model with *Phaser* of *Pheni*x software suite ^61^. The electron density block of TNA was clearly shown in both 2F_o_–F_c_ and F_o_–F_c_ maps. Subsequently, TNA structure was manually constructed, followed by several rounds of refinement using *Phenix.refine* ^58^, with manual fitting and rebuilding using *COOT* ^60^. We also determined the structure of phloroglucinol-soaked crystals at a resolution of 1.4 Å via the MR method using the refined EHEP structure as a search model with *Phaser*, but no electron density block of phloroglucinol was obtained. Therefore, we referred to this structure as the apo form (apo structure2). The final refinement statistics and geometry are shown in Table 1.

We determined the structure of *aku*BGL via the MR method using *Phaser* of *Phenix* software suite ^61^. We used one GH domain (86–505 aa) of β-klotho (PDB entry: 5VAN) ^62^ as the search model. This GH domain of β-klotho shares 30% sequence identity with *aku*BGL. Four GH domains of two molecules in an asymmetric unit were found and subsequently rebuilt with *Phenix_autobuild* of *Phenix* software suite ^58^. Finally, refinement of *aku*BGL structure was performed as described for EHEP.

### Docking studies of *aku*BGL with phlorotanins and laminarins

We used Schrodinger Maestro program for performing docking studies ^63^. First, we superimposed the structure of *Os*BGL mutant complexed with cellotetraose (PDB ID 4QLK ^64^) to that of *aku*BGL GH1D2 to define the ligand position in the ligand-binding cavity. Then, we modified the structure of the *aku*BGL GH1D2 using wizard module to remove water molecules and add hydrogen atoms for docking. The 2D structures of the inhibition ligands, including TNA, phloroglucinol, and eckol, were downloaded from Pubchem ^65^ and further converted to 3D structures using the LigPrep module of Schrodinger Maestro program. The structure of the substate laminaritetraose was extracted from the *Zobellia galactanivorans β*-glucanase–laminaritetraose complex structure (PDB ID: 4BOW) ^66^. Then, a receptor grid was constructed in the center of the ligand-binding cavity. We performed docking using the Glide standard precision mode without any constraints. The optimal binding pose was determined using the lowest Glide score, and docked structures were analyzed using PyMol.

### Data Availability

The atomic coordinates were deposited in the PDB with the accession codes as follows: EHEP with 1.15 Å resolution (8IN3), EHEP with 1.4 Å resolution (8IN4), EHEP complexed with tannic acid (8IN6), *aku*BGL(8IN1).

## Acknowledgments

This work was supported in part by Grant-in-Aid for Scientific Research (B) (Grant Number 21H01754 To M. Y) and Platform Project for Supporting Drug Discovery and Life Science Research (Basis for Supporting Innovative Drug Discovery and Life Science Research (BINDS)) from Japan Agency for Medical Research and Development (AMED) under Grant Number JP18am0101071 and JP19am0101083. We are grateful to the Photon Factor and SPring-8 (No. 2017B2545, 2017A2551, 2018B2538) for beam time and the beamline staff for their assistance for data collection.

## Supporting information

**Fig. S1.**
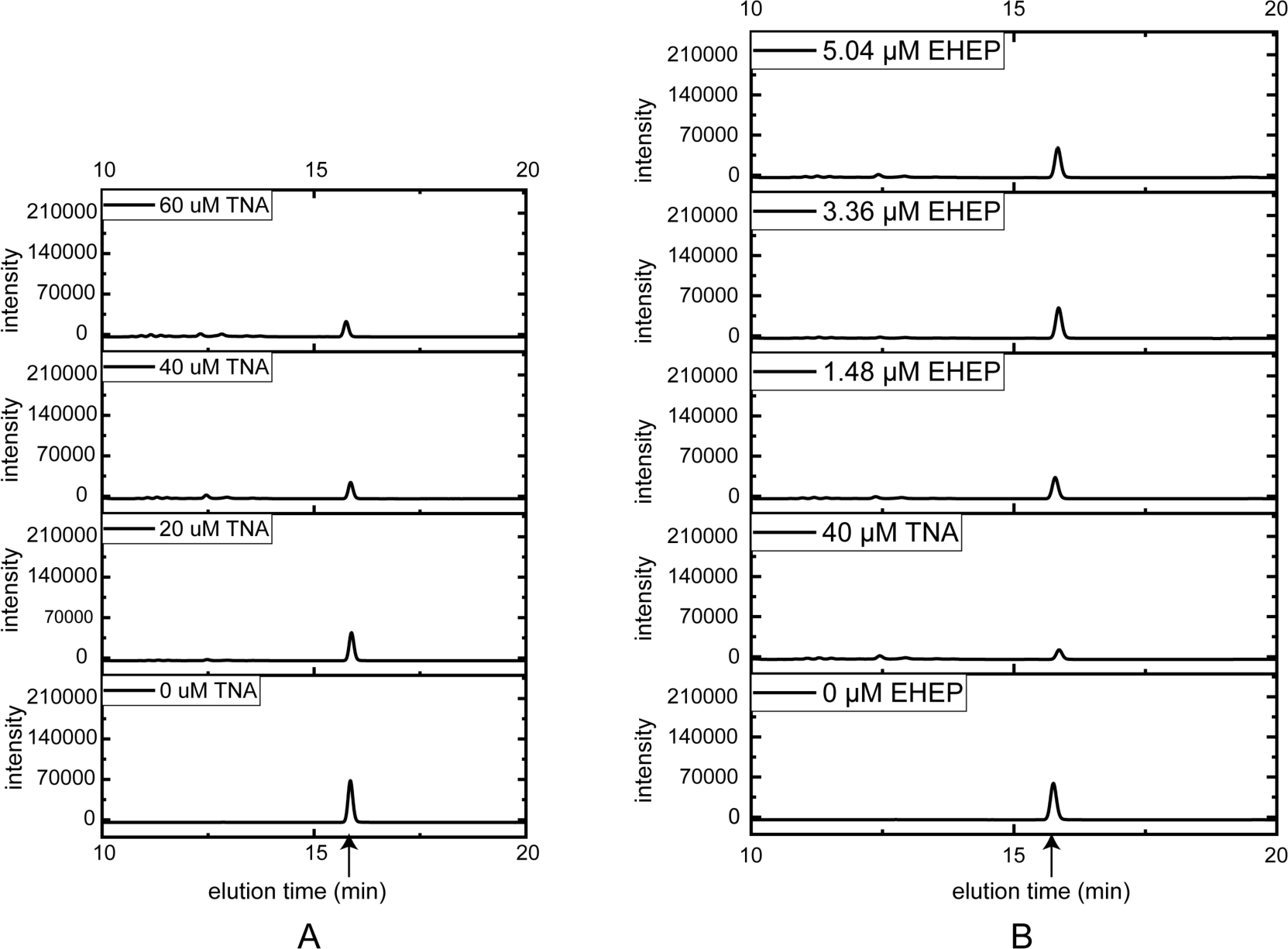
HPLC profiles of *aku*BGL activity toward ortho-Nitrophenyl-β-galactoside. (A) TNA inhibition *aku*BGL by increasing the concentration of TNA with 0.049 μM *aku*BGL and 2.5 mM ortho-Nitrophenyl-β-galactoside. (B) EHEP protects *aku*BGL from TNA inhibition by increasing the concentration of EHEP. The product peak was marked by a black arrow.

**Figure. S2.**
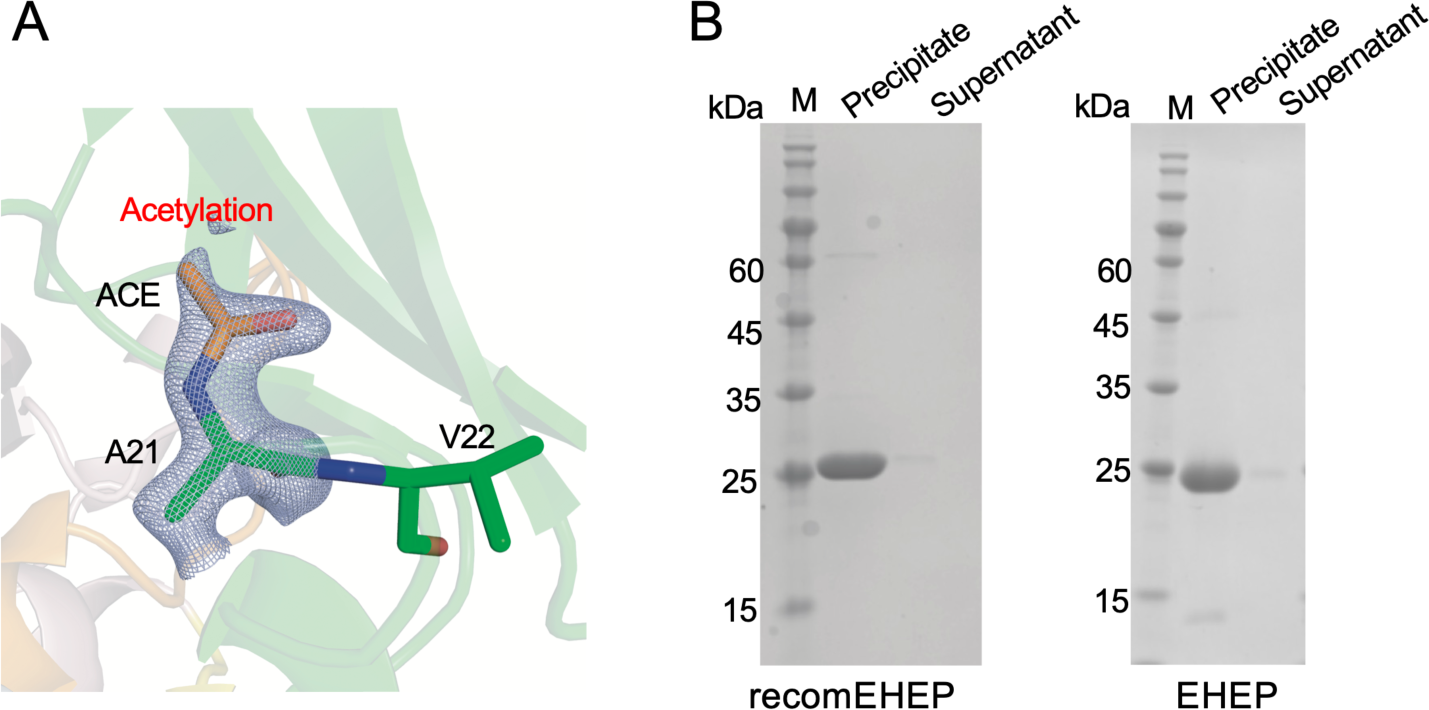
(A) Acetylation modification on the N-terminal residue A21. The structure is shown as sticks with an omitted map of the acetylation of A21 at a 3.3 σ level (blue-white). (B) TNA binding activity of recomEHEP and EHEP. SDS–PAGE was run using a mixture of recomEHEP/EHEP and TNA.

**Figure S3.**
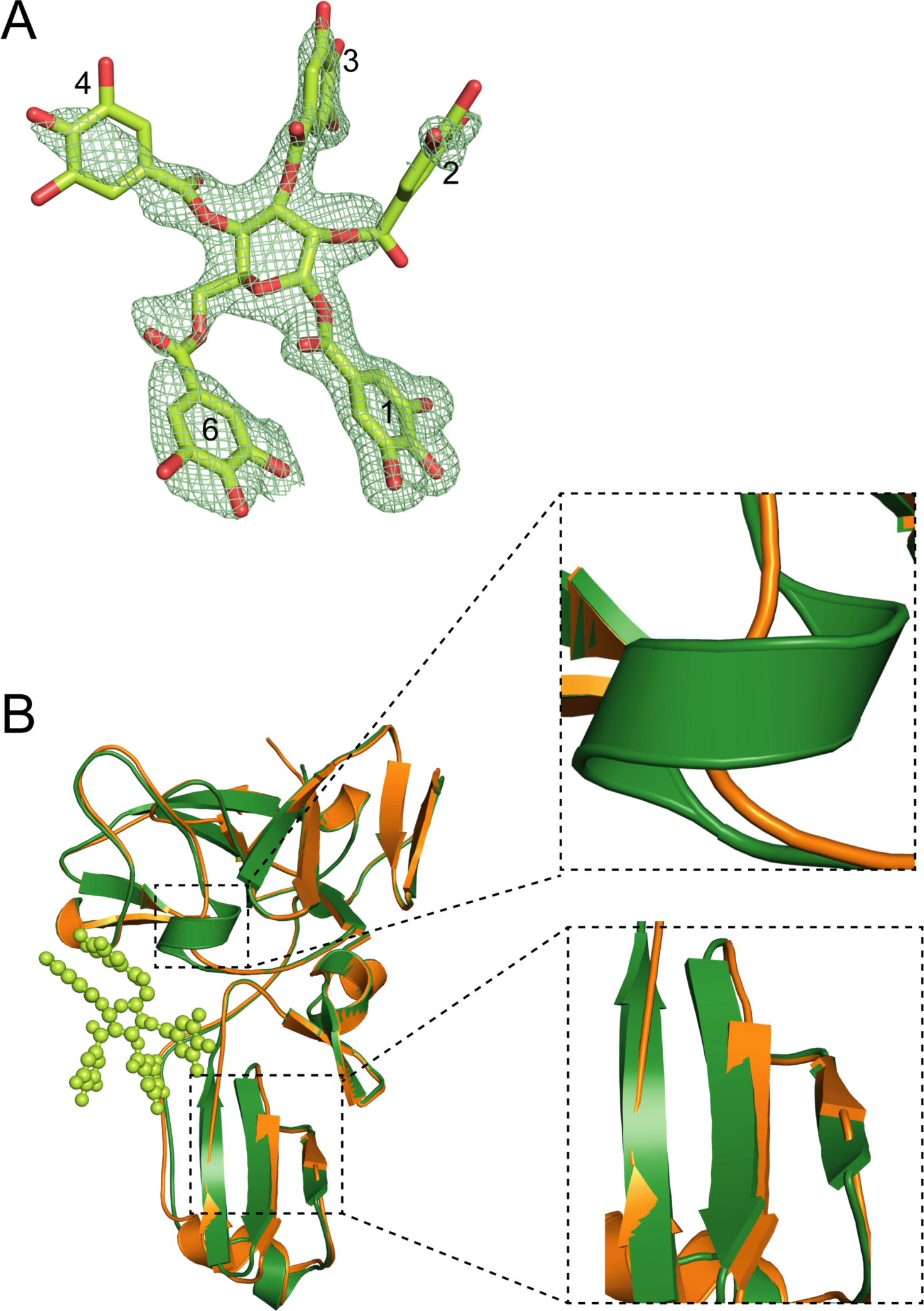
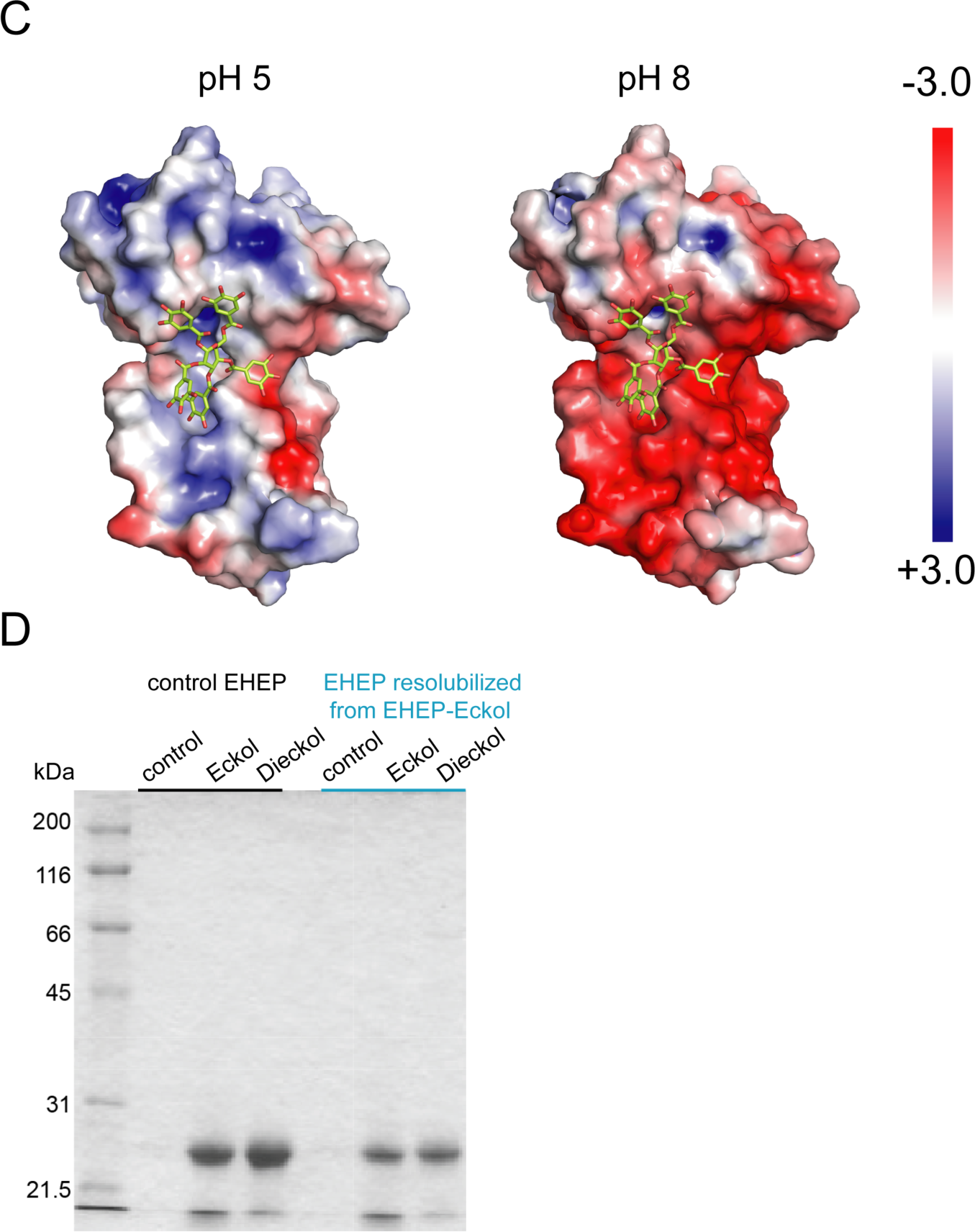
EHEP–TNA structure and structural comparisons. (A) TNA structure (stick model) in EHEP–TNA with an omitted map countered at the 2.0 σ level. The O and C atoms are colored red and lemon, respectively. (B) Structure superposition of apo EHEP (green) with EHEP–TNA (orange). The conformational changes are marked by black dotted boxes and zoomed on the right. (C) Electrostatic potential of the EHEP surface in the EHEP–TNA complex at different pH values, which were calculated using APBS-PDB2PQR software suite. (D) Activity of the resolubilized EHEP. The EHEP–eckol precipitate was resolubilized in Tris-HCl (pH 8.0) buffer and then the supernatant was purified using a Sephacryl S-100 HR column (2.0 ×110 cm) to measure eckol/dieckol-binding activity.

**Figure S4.**
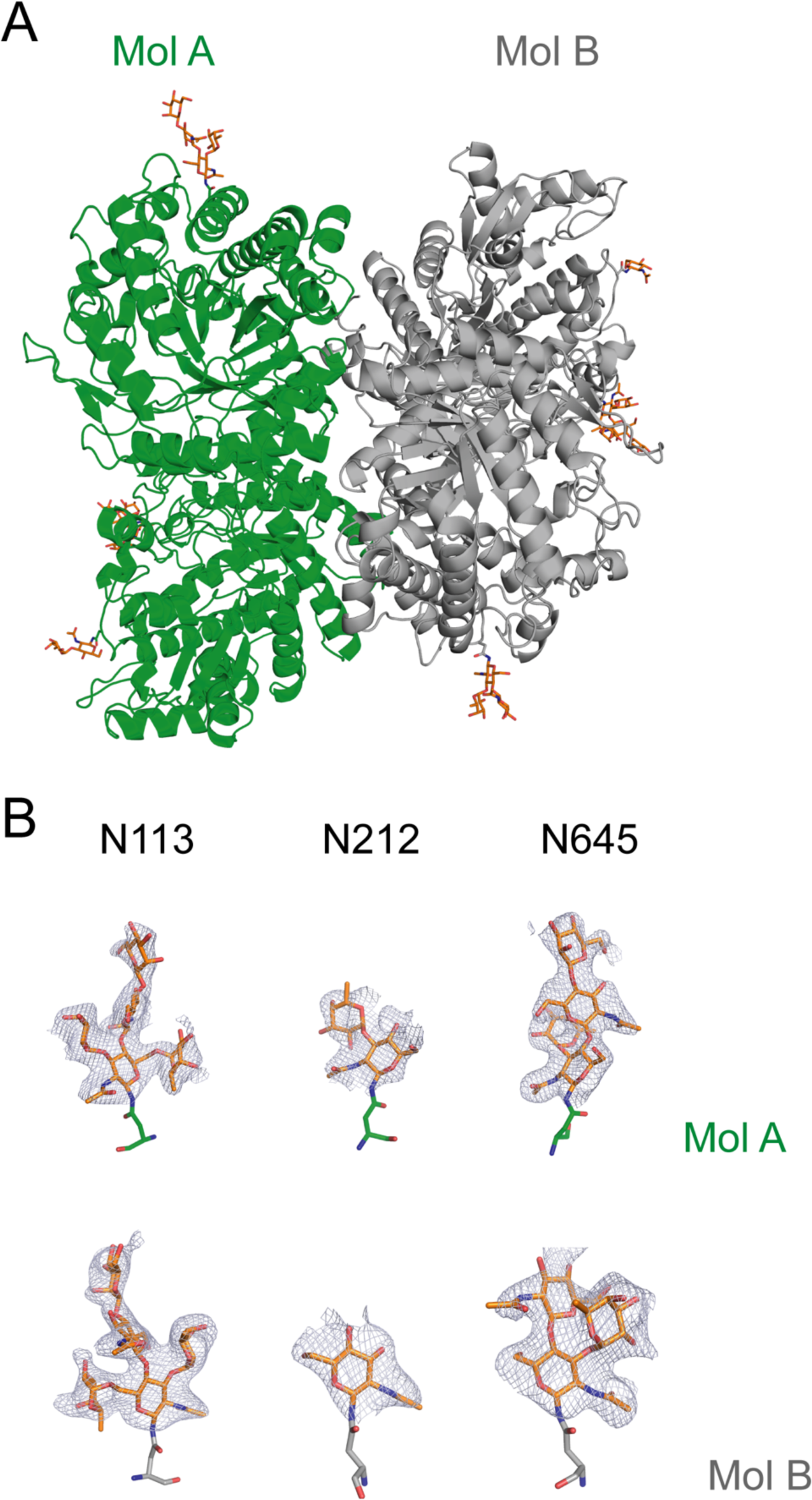
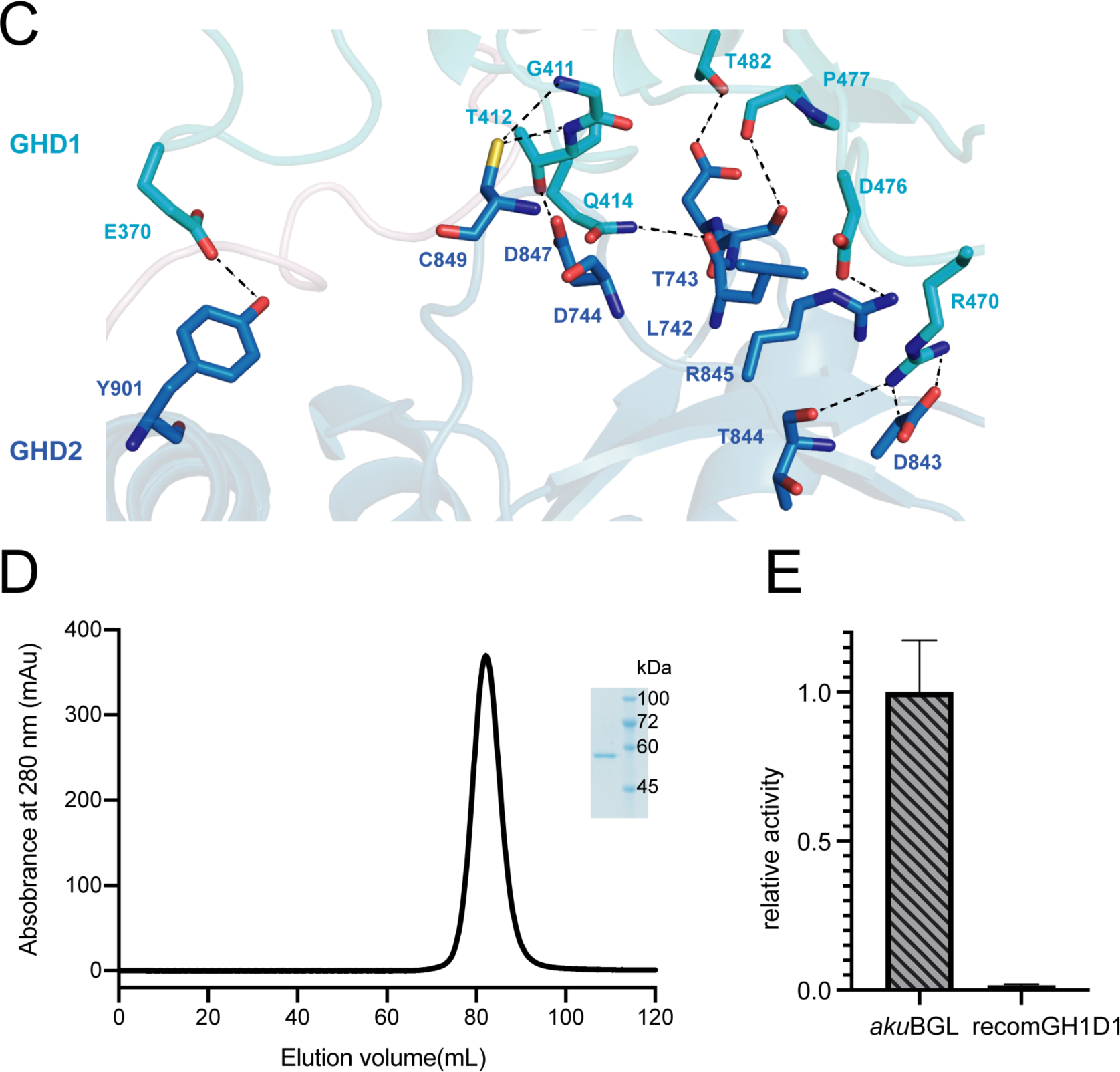
*aku*BGL structure. (A) Two *aku*BGL molecules in the asymmetric unit, colored green and grey, respectively. The glycosylation sites are shown as orange sticks, with O and N atoms in red and blue, respectively. (B) The glycosylation chains with omitted density maps are countered at the 2.0 σ level. (C) The interface between the GH1D1 (cyan) and GH1D2 (blue). The key interactions between the two domains are shown as black dashed lines. (D) Size exclusion chromatogram of the purified GH1D1 domain. The inset shows the SDS–PAGE analysis of the GH1D1 domain. (E) Galactoside hydrolytic activity of the recomGH1D1 toward ortho-Nitrophenyl-β-galactoside. The average and standard deviation of the relative activity were estimated from three independent replicates (N = 3).

**Figure S5.**
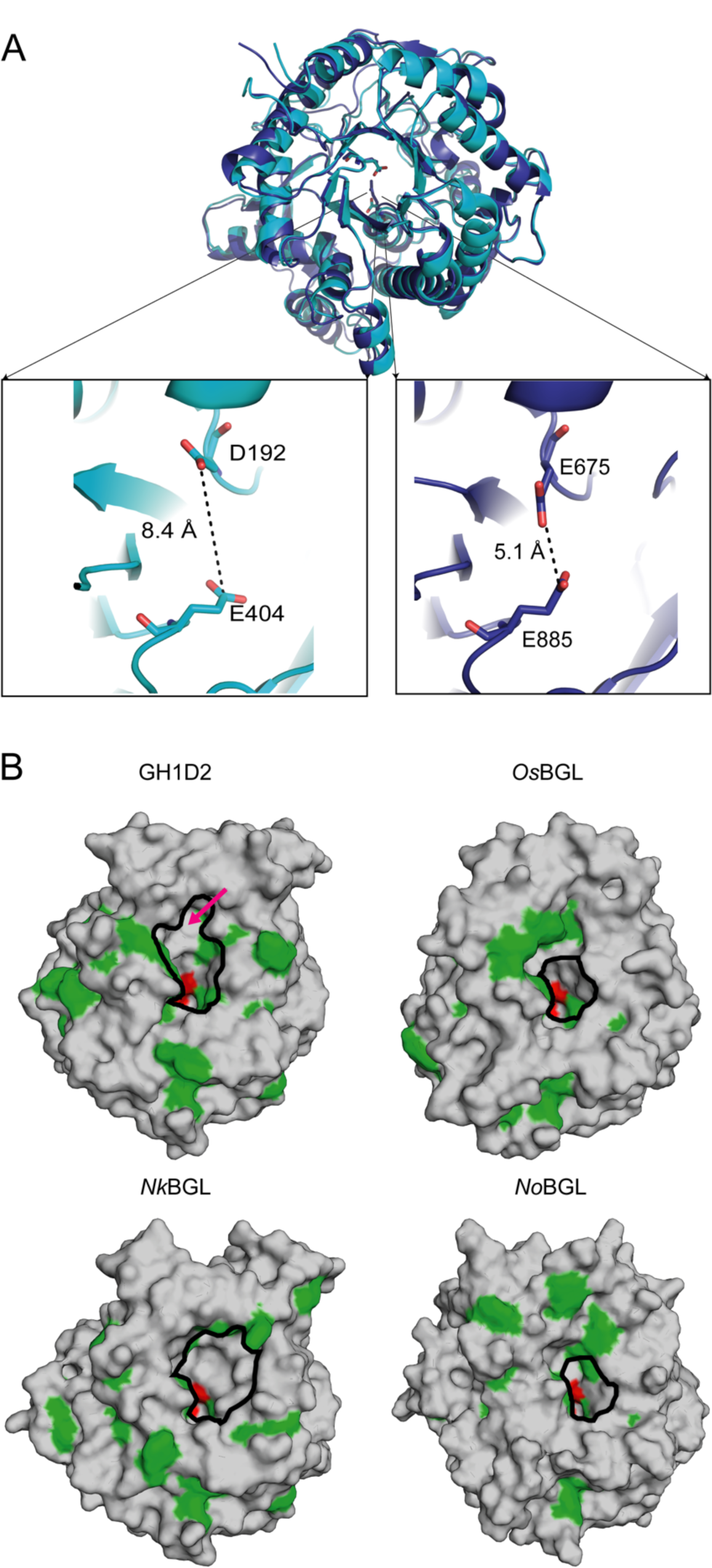
Structural comparison of BGLs. (A) Structure superposition of GH1D1(cyan) and GH1D2 (blue). The enlarged picture shows the distance of conceivable catalytic residues in GH1D1 and GH1D2. (B) Surface representations of GH1D2, *Nk*BGL(PDB ID 3VIH) ^32^, *Os*BGL(PDB ID 2RGL) ^35^, and *No*BGL (PDB ID 5YJ7) ^33^ with the aromatic and catalytic residues colored green and red, respectively. The red arrow indicates the location of the auxiliary site of GH1D2. The active pockets are highlighted by a black circle on each surface.

**Figure S6.**
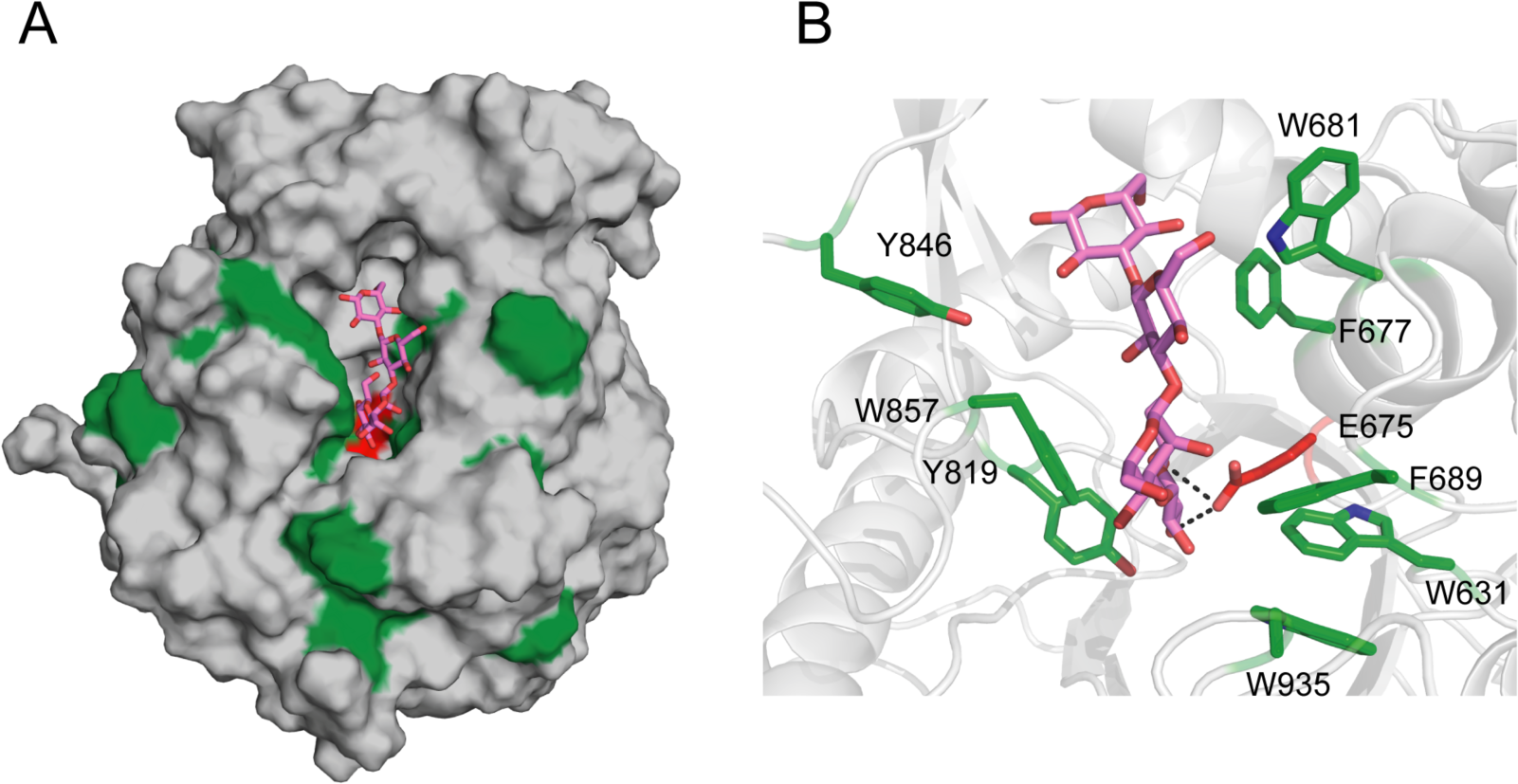
The model of GH1D2 docking with the substrate laminaritetraose. (A) Overall model of GH1D2-laminaritetraose. GH1D2 is shown as a grey surface representation and laminaritetraose as marine sticks. The aromatic and catalytic residues of GH1D2 are colored green and red, respectively. (B) The closeup review of interaction between laminatrtetraose and GH1D2 with key residues showing in sticks.

**Figure S7.**
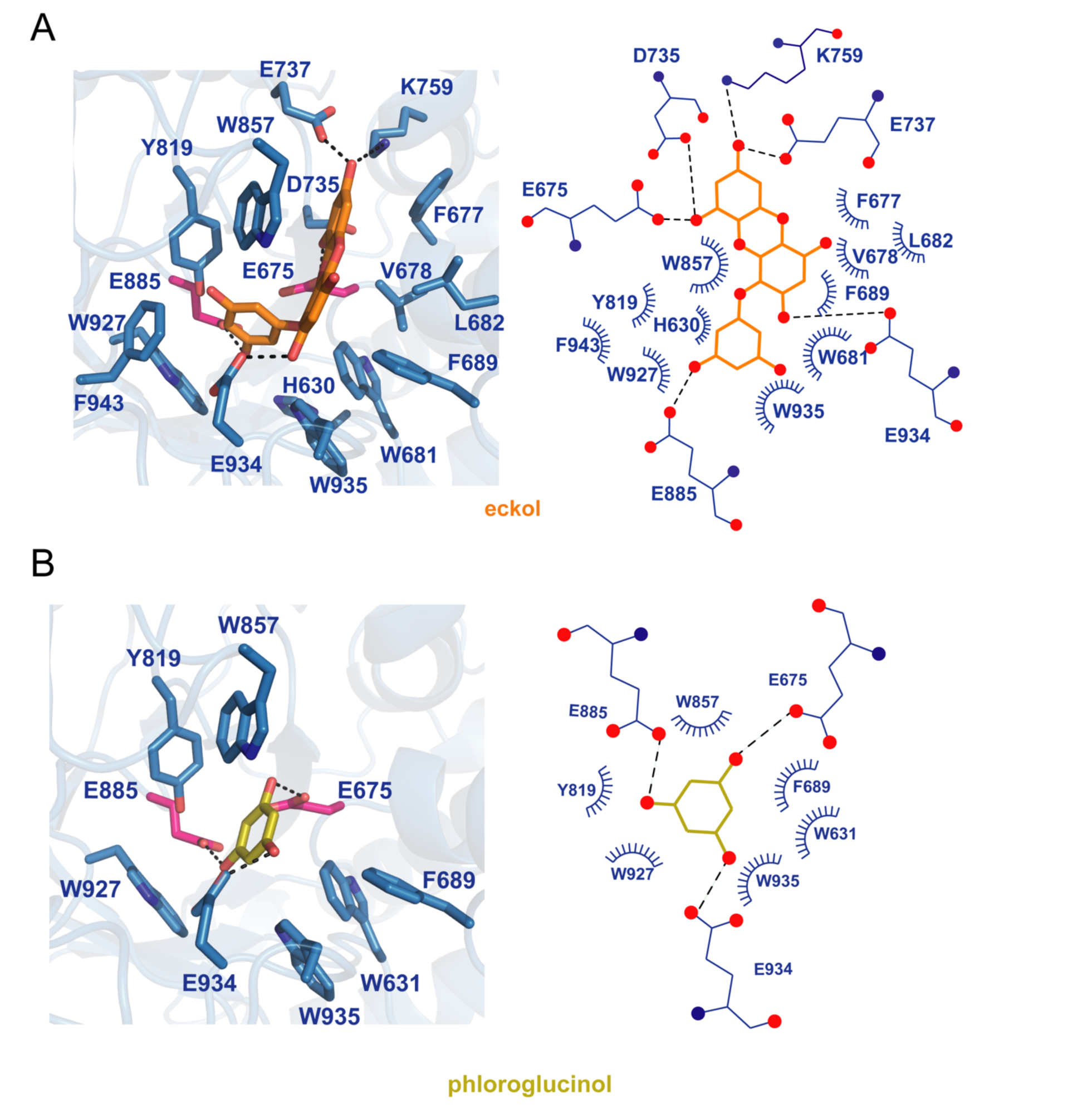
The models of GH1D2 docking. (A) With eckol, and (B) With phloroglucinol. The left panel shows the 3D structures, and the right panel shows the 2D diagrams. The C, N, and O atoms of residues are colored light blue, dark blue, and red, respectively. The C atoms of eckol and phloroglucinol are shown in orange and yellow, respectively.

## Notes

### Competing Interest Statement

The authors have declared no competing interest.

### Summary of Updates

1. Method updated including: added the recomGH1D1 expression and purification;added the activity assay of recomGH1D1; added N-ter sequencing of akuBGL; updated the resolubiliztion of EHEP and activity. 2 Supplementary updated including: moved fig.3 to Fig. S2; added Fig. S1 to show the HPLC result.

## References

1. Becklin, K.M. A Coevolutionary Arms Race: Understanding Plant-Herbivore Interactions. The American Biology Teacher 70, 288–292 (2008).

2. Jormalainen, V. & Honkanen, T. in Algal Chemical Ecology 57–89 (2008).

3. Enquist-Newman, M. et al. Efficient ethanol production from brown macroalgae sugars by a synthetic yeast platform. Nature 505, 239–243 (2014).

4. Tsuji, A., Kuwamura, S., Shirai, A. & Yuasa, K. Identification and Characterization of a 25 kDa Protein That Is Indispensable for the Efficient Saccharification of Eisenia bicyclis in the Digestive Fluid of Aplysia kurodai. PLoS One 12, e0170669 (2017).

5. Erb, M. & Reymond, P. Molecular Interactions Between Plants and Insect Herbivores. Annu Rev Plant Biol 70, 527–557 (2019).

6. Mendez-Liter, J.A., de Eugenio, L.I., Prieto, A. & Martinez, M.J. The beta-glucosidase secreted by Talaromyces amestolkiae under carbon starvation: a versatile catalyst for biofuel production from plant and algal biomass. Biotechnol Biofuels 11, 123 (2018).

7. Nakajima, M., Yamashita, T., Takahashi, M., Nakano, Y. & Takeda, T. Identification, cloning, and characterization of beta-glucosidase from Ustilago esculenta. Appl Microbiol Biotechnol 93, 1989–1998 (2012).

8. Wang, Z. et al. A novel Vibrio beta-glucosidase (LamN) that hydrolyzes the algal storage polysaccharide laminarin. FEMS Microbiol Ecol 91, fiv087 (2015).

9. Kim, D.H., Kim, D.H., Lee, S.H. & Kim, K.H. A novel beta-glucosidase from Saccharophagus degradans 2-40(T) for the efficient hydrolysis of laminarin from brown macroalgae. Biotechnol Biofuels 11, 64 (2018).

10. Mohsin, I., Poudel, N., Li, D.C. & Papageorgiou, A.C. Crystal Structure of a GH3 beta-Glucosidase from the Thermophilic Fungus Chaetomium thermophilum. Int J Mol Sci 20 (2019).

11. Hegedus, D., Erlandson, M., Gillott, C. & Toprak, U. New Insights into Peritrophic Matrix Synthesis, Architecture, and Function. Annual Review of Entomology 54, 285–302 (2009).

12. Devenport, M. et al. Identification of the Aedes aegypti peritrophic matrix protein AeIMUCI as a heme-binding protein. Biochemistry 45, 9540–9549 (2006).

13. Cassani, L. et al. Seaweed-based natural ingredients: Stability of phlorotannins during extraction, storage, passage through the gastrointestinal tract and potential incorporation into functional foods. Food Res Int 137, 109676 (2020).

14. Imbs, T.I. & Zvyagintseva, T.N. Phlorotannins are Polyphenolic Metabolites of Brown Algae. Russian Journal of Marine Biology 44, 263–273 (2018).

15. Holm, L. & Rosenstrom, P. Dali server: conservation mapping in 3D. Nucleic Acids Res 38, W545–549 (2010).

16. Fadel, F. et al. X-Ray Crystal Structure of the Full Length Human Chitotriosidase (CHIT1) Reveals Features of Its Chitin Binding Domain. PLoS One 11, e0154190 (2016).

17. Kohler, A.C. et al. Structural Analysis of an Avr4 Effector Ortholog Offers Insight into Chitin Binding and Recognition by the Cf-4 Receptor. Plant Cell 28, 1945–1965 (2016).

18. Hurlburt, N.K., Chen, L.H., Stergiopoulos, I. & Fisher, A.J. Structure of the Cladosporium fulvum Avr4 effector in complex with (GlcNAc)6 reveals the ligand-binding mechanism and uncouples its intrinsic function from recognition by the Cf-4 resistance protein. PLoS Pathog 14, e1007263 (2018).

19. Mueller, G.A. et al. Serological, genomic and structural analyses of the major mite allergen Der p 23. Clin Exp Allergy 46, 365–376 (2016).

20. Suetake, T. et al. Chitin-binding proteins in invertebrates and plants comprise a common chitin-binding structural motif. J Biol Chem 275, 17929–17932 (2000).

21. Madland, E., Crasson, O., Vandevenne, M., Sorlie, M. & Aachmann, F.L. NMR and Fluorescence Spectroscopies Reveal the Preorganized Binding Site in Family 14 Carbohydrate-Binding Module from Human Chitotriosidase. ACS Omega 4, 21975–21984 (2019).

22. Sun, X. et al. Crystallographic analysis of Eisenia hydrolysis-enhancing protein using a long wavelength for native-SAD phasing. Acta Crystallographica Section F-Structural Biology Communications 76, 20–24 (2020).

23. Silva, R.D. & Martinho, R.G. Developmental roles of protein N-terminal acetylation. Proteomics 15, 2402–2409 (2015).

24. Hollebeke, J., Van Damme, P. & Gevaert, K. N-terminal acetylation and other functions of Nalpha-acetyltransferases. Biol Chem 393, 291–298 (2012).

25. Lange, P.F. & Overall, C.M. TopFIND, a knowledgebase linking protein termini with function. Nat Methods 8, 703–704 (2011).

26. Jie Fu, J.x., Zhang, Y. & Lu, X. A Greener Process for Gallic Acid Production from Tannic Acid Hydrolysis with Hydrochloric Acid. Asian Journal of Chemistry 27, 3328–3332 (2015).

27. Luo, Q. et al. A novel green process for tannic acid hydrolysis using an internally sulfonated hollow polystyrene sphere as catalyst. RSC Advances 8, 17151–17158 (2018).

28. Amore, A. et al. Distinct roles of N- and O-glycans in cellulase activity and stability. Proc Natl Acad Sci U S A 114, 13667–13672 (2017).

29. Han, C. et al. Improvement of the catalytic activity and thermostability of a hyperthermostable endoglucanase by optimizing N-glycosylation sites. Biotechnol Biofuels 13, 30 (2020).

30. Wei, W. et al. N-glycosylation affects the proper folding, enzymatic characteristics and production of a fungal beta-glucosidas. Biotechnol Bioeng 110, 3075–3084 (2013).

31. Hayashi, Y. et al. Klotho-related protein is a novel cytosolic neutral beta-glycosylceramidase. J Biol Chem 282, 30889–30900 (2007).

32. Jeng, W.Y. et al. High-resolution structures of Neotermes koshunensis beta-glucosidase mutants provide insights into the catalytic mechanism and the synthesis of glucoconjugates. Acta Crystallogr D Biol Crystallogr 68, 829–838 (2012).

33. Dong, S. et al. Structural insight into a GH1 beta-glucosidase from the oleaginous microalga, Nannochloropsis oceanica. Int J Biol Macromol 170, 196–206 (2021).

34. Tamaki, F.K. et al. Using the Amino Acid Network to Modulate the Hydrolytic Activity of beta-Glycosidases. PLoS One 11, e0167978 (2016).

35. Chuenchor, W. et al. Structural insights into rice BGlu1 beta-glucosidase oligosaccharide hydrolysis and transglycosylation. J Mol Biol 377, 1200–1215 (2008).

36. Opassiri, R. et al. Characterization of a rice β-glucosidase highly expressed in flower and germinating shoot. Plant Science 165, 627–638 (2003).

37. Hudson, K.L. et al. Carbohydrate-Aromatic Interactions in Proteins. J Am Chem Soc 137, 15152–15160 (2015).

38. Ni, J., Tokuda, G., Takehara, M. & Watanabe, H. Heterologous expression and enzymatic characterization of .BETA.-glucosidase from the drywood-eating termite, Neotermes koshunensis. Applied Entomology and Zoology 42, 457–463 (2007).

39. Tsuji, A., Tominaga, K., Nishiyama, N. & Yuasa, K. Comprehensive enzymatic analysis of the cellulolytic system in digestive fluid of the Sea Hare Aplysia kurodai. Efficient glucose release from sea lettuce by synergistic action of 45 kDa endoglucanase and 210 kDa ss-glucosidase. PLoS One 8, e65418 (2013).

40. Jung, H.A., Oh, S.H. & Choi, J.S. Molecular docking studies of phlorotannins from Eisenia bicyclis with BACE1 inhibitory activity. Bioorg Med Chem Lett 20, 3211–3215 (2010).

41. Amsler, C.D. & Fairhead, V.A. in Advances in Botanical Research, Vol 43: Incorporating Advances in Plant Pathology, Vol. 43. (ed. J.A. Callow) 1-91 (2006).

42. Sakamoto, K., Uji, S., Kurokawa, T. & Toyohara, H. Molecular cloning of endogenous beta-glucosidase from common Japanese brackish water clam Corbicula japonica. Gene 435, 72–79 (2009).

43. Sabbadin, F. et al. Uncovering the molecular mechanisms of lignocellulose digestion in shipworms. Biotechnol Biofuels 11, 59 (2018).

44. Barbehenn, R.V. & Peter Constabel, C. Tannins in plant-herbivore interactions. Phytochemistry 72, 1551–1565 (2011).

45. Marsh, K.J. et al. New approaches to tannin analysis of leaves can be used to explain in vitro biological activities associated with herbivore defence. New Phytol 225, 488–498 (2020).

46. Shimada, T. Salivary proteins as a defense against dietary tannins. J Chem Ecol 32, 1149–1163 (2006).

47. De Smet, K. & Contreras, R. Human Antimicrobial Peptides: Defensins, Cathelicidins and Histatins. Biotechnology Letters 27, 1337–1347 (2005).

48. War, A.R. et al. in Co-Evolution of Secondary Metabolites 795–822 (2020).

49. Lin, D. et al. The effect of ionic strength and pH on the stability of tannic acid-facilitated carbon nanotube suspensions. Carbon 47, 2875–2882 (2009).

50. Ge, D., Yuan, H., Shen, Y., Zhang, W. & Zhu, N. Improved sludge dewaterability by tannic acid conditioning: Temperature, thermodynamics and mechanism studies. Chemosphere 230, 14–23 (2019).

51. Yi, Z. et al. Green, effective chemical route for the synthesis of silver nanoplates in tannic acid aqueous solution. Colloids and Surfaces A: Physicochemical and Engineering Aspects 392, 131–136 (2011).

52. Dultz, S., Mikutta, R., Kara, S.N.M., Woche, S.K. & Guggenberger, G. Effects of solution chemistry on conformation of self-aggregated tannic acid revealed by laser light scattering. Sci Total Environ 754, 142119 (2021).

53. Han, Y. et al. Polyphenol-Mediated Assembly of Proteins for Engineering Functional Materials. Angew Chem Int Ed Engl 59, 15618–15625 (2020).

54. Lemke, T., Stingl, U., Egert, M., Friedrich, M.W. & Brune, A. Physicochemical conditions and microbial activities in the highly alkaline gut of the humus-feeding larva of Pachnoda ephippiata (Coleoptera: Scarabaeidae). Appl Environ Microbiol 69, 6650–6658 (2003).

55. Yao, M. & Li, L. (NATIONAL UNIVERSITY CORPORATION HOKKAIDO UNIVERSITY 2022).

56. Kabsch, W. Xds. Acta Crystallogr D Biol Crystallogr 66, 125–132 (2010).

57. Yu, J., Shinoda, A., Kato, K., Tanaka, I. & Yao, M. A solution-free crystal-mounting platform for native SAD. Acta Crystallogr D Struct Biol 76, 938–945 (2020).

58. Adams, P.D. et al. PHENIX: a comprehensive Python-based system for macromolecular structure solution. Acta Crystallogr D Biol Crystallogr 66, 213–221 (2010).

59. Afonine, P.V. et al. Towards automated crystallographic structure refinement with phenix.refine. Acta Crystallogr D Biol Crystallogr 68, 352–367 (2012).

60. Emsley, P. & Cowtan, K. Coot: model-building tools for molecular graphics. Acta Crystallogr D Biol Crystallogr 60, 2126–2132 (2004).

61. McCoy, A.J., et al. Phaser crystallographic software. J Appl Crystallogr 40, 658–674 (2007).

62. Lee, S. et al. Structures of beta-klotho reveal a ’zip code’-like mechanism for endocrine FGF signalling. Nature 553, 501–505 (2018).

63. Sastry, G.M., Adzhigirey, M., Day, T., Annabhimoju, R. & Sherman, W. Protein and ligand preparation: parameters, protocols, and influence on virtual screening enrichments. Journal of Computer-Aided Molecular Design 27, 221–234 (2013).

64. Pengthaisong, S. & Ketudat Cairns, J.R. Effects of active site cleft residues on oligosaccharide binding, hydrolysis, and glycosynthase activities of rice BGlu1 and its mutants. Protein Sci 23, 1738–1752 (2014).

65. Wang, Y.L. et al. PubChem: a public information system for analyzing bioactivities of small molecules. Nucleic Acids Research 37, W623–W633 (2009).

66. Labourel, A. et al. The beta-glucanase ZgLamA from Zobellia galactanivorans evolved a bent active site adapted for efficient degradation of algal laminarin. J Biol Chem 289, 2027–2042 (2014).

67. Hakulinen, N., Paavilainen, S., Korpela, T. & Rouvinen, J. The crystal structure of beta-glucosidase from Bacillus circulans sp. alkalophilus: ability to form long polymeric assemblies. J Struct Biol 129, 69–79 (2000).

